# ELOF1 is a core component of the promoter-proximal paused RNA polymerase II complex

**DOI:** 10.64898/2026.05.26.728003

**Authors:** Roberto J. Vazquez Nunez, Annkatrin Bressin, Zhihao Shao, Andreas Mayer, Seychelle M. Vos

**Author notes:** Correspondence should be addressed to S.M.V.

## Abstract

Metazoan RNA polymerase II is prone to pause in the promoter-proximal region of genes. This pausing involves DRB Sensitivity-Inducing Factor (DSIF) and Negative Elongation Factor (NELF). DSIF and NELF, however, are insufficient to fully reconstitute the degree of promoter-proximal pausing observed in cells. Their negative effects are counteracted by transcription initiation factor TFIIF or by physiological nucleotide concentrations, indicating an additional factor is required. Here, we report that Elongation Factor Homolog 1 (ELOF1) is this missing factor. ELOF1 is enriched in the promoter-proximal region of genes, and its rapid degradation reduces pause duration in cells. In reconstituted assays, ELOF1 potently enhances pausing induced by DSIF and NELF at physiological nucleotide concentrations. Cryo-EM structures reveal that DSIF-NELF-ELOF1 sterically clashes with the position of TFIIF on RNA polymerase II. Accordingly, RNA polymerase II-DSIF-NELF-ELOF1, but not RNA polymerase II-DSIF-NELF, counteracts the positive effects of TFIIF. Our results establish ELOF1 as a core component of promoter-proximal paused RNA polymerase II.

## Introduction

Metazoan RNA polymerase II is responsible for transcribing most protein coding mRNAs and is prone to pause after synthesizing the first 20-100 base pairs (bp) of a gene^1– 4^. These promoter-proximal pausing events serve a critical regulatory checkpoint to determine if transcription proceeds into productive elongation or is prematurely terminated at the pause site^3,5^. Promoter-proximal pausing involves protein complexes DRB Sensitivity-Inducing Factor (DSIF) and Negative Elongation Factor (NELF)^6–11^. The RNA polymerase II-DSIF-NELF complex can be either activated and released into productive and processive elongation or prematurely terminated^3,5,12^. For activation, the kinase positive-transcription elongation factor (P-TEF) b phosphorylates RNA polymerase II, DSIF, and NELF, enabling the association of elongation factors like SPT6 and PAF1c and displacement of NELF on RNA polymerase II^13–20^. Premature termination is induced by factors including Integrator and the E3 ubiquitin ligase ARMC5^12,21–29^.

Reconstituted systems incompletely recapitulate the degree of promoter-proximal pausing observed in cells. Specifically, limiting nucleotide concentrations are required to recapitulate the pausing levels observed in cells, and transcription initiation factor TFIIF can counteract the negative effects of DSIF and NELF^30,31^. Prior work with partially purified complexes showed that addition of nuclear extract could make previously sensitive RNA polymerase II-DSIF-NELF complexes resistant to TFIIF^31^. This indicated the presence of an extract specific “TFIIF resistance factor”^31^. Importantly, this resistance factor was assigned roles in both promoter-proximal pausing and transcription elongation^31^. The resistance factor was initially identified as GDOWN1^32,33^. Subsequent studies, however, found that GDOWN1 primarily resides in the cytoplasm, and appears to mostly associate with RNA polymerase II during mitosis^34,35^. Thus, it has remained unclear which factor(s) is responsible for endowing RNA polymerase II-DSIF-NELF complexes with TFIIF resistance and the ability to pause at physiological nucleotide concentrations.

Here we show that Elongation Factor Homolog 1 (ELOF1) enhances promoter-proximal pausing and can act as the “TFIIF resistance factor” with DSIF-NELF. Human ELOF1 is an 83 amino acid protein broadly conserved in archaea and eukaryotes, has well defined roles as a transcription elongation factor, is essential for mouse embryonic development, and is primarily localized to the nucleus^36–40^. During transcription elongation, ELOF1 directly associates with DSIF and appears to stabilize the jaw and clamp domains of RNA polymerase II^41–43^. Recent work has shown that ELOF1 facilitates transcription-coupled nucleotide excision repair (TC-NER), class switch recombination (CSR), and somatic hypermutation (SHM)^44,45^. These studies have collectively hinted at a potential role of ELOF1 in promoter-proximal pausing^38,39,44,45^. For instance, complete knockout of ELOF1 showed decreased NELF association with RNA polymerase II, and ELOF1 was found to be enriched within promoter-proximal regions of genes, suggesting that it associates with RNA polymerase II shortly after promoter escape^38,39,44^. Using cell based, biochemical, and structural approaches, we show that ELOF1 enhances promoter-proximal pausing in cells, allows for the reconstitution of RNA polymerase II pausing behavior under physiological conditions, and is the long sought after “TFIIF resistance factor”.

## Results

### ELOF1 is enriched in the promoter-proximal region of genes

We first assessed whether ELOF1 is involved in promoter-proximal pausing in cells utilizing two previously established human cell lines where ELOF1 was either fused to a degron tag to allow for its selective and rapid depletion (ELOF1-degron) or stably knocked out (ΔELOF1)^39,44^. The rapid and selective degradation of proteins in cells can overcome some of the limitations associated with complete gene knockouts^46^. We thus first utilized human RASH-1C cells encoding ELOF1-HA-FKPB12^F36V^-mScarlet (ELOF1-degron)^44^ (**Supplementary Fig. 1A**). Rapid degradation of ELOF1 is observed upon treatment of cells with dTAGV-1 but not with control compound dTAG-NEG (**Supplementary Fig. 1B**). ChIP-nexus using anti-HA antibodies showed that ELOF1-degron occupancy was enriched within the promoter-proximal regions of genes (**Fig. 1A**). This observation is consistent with previous CUT&RUN-seq data performed in the same cell line^44^.

**Figure 1.**
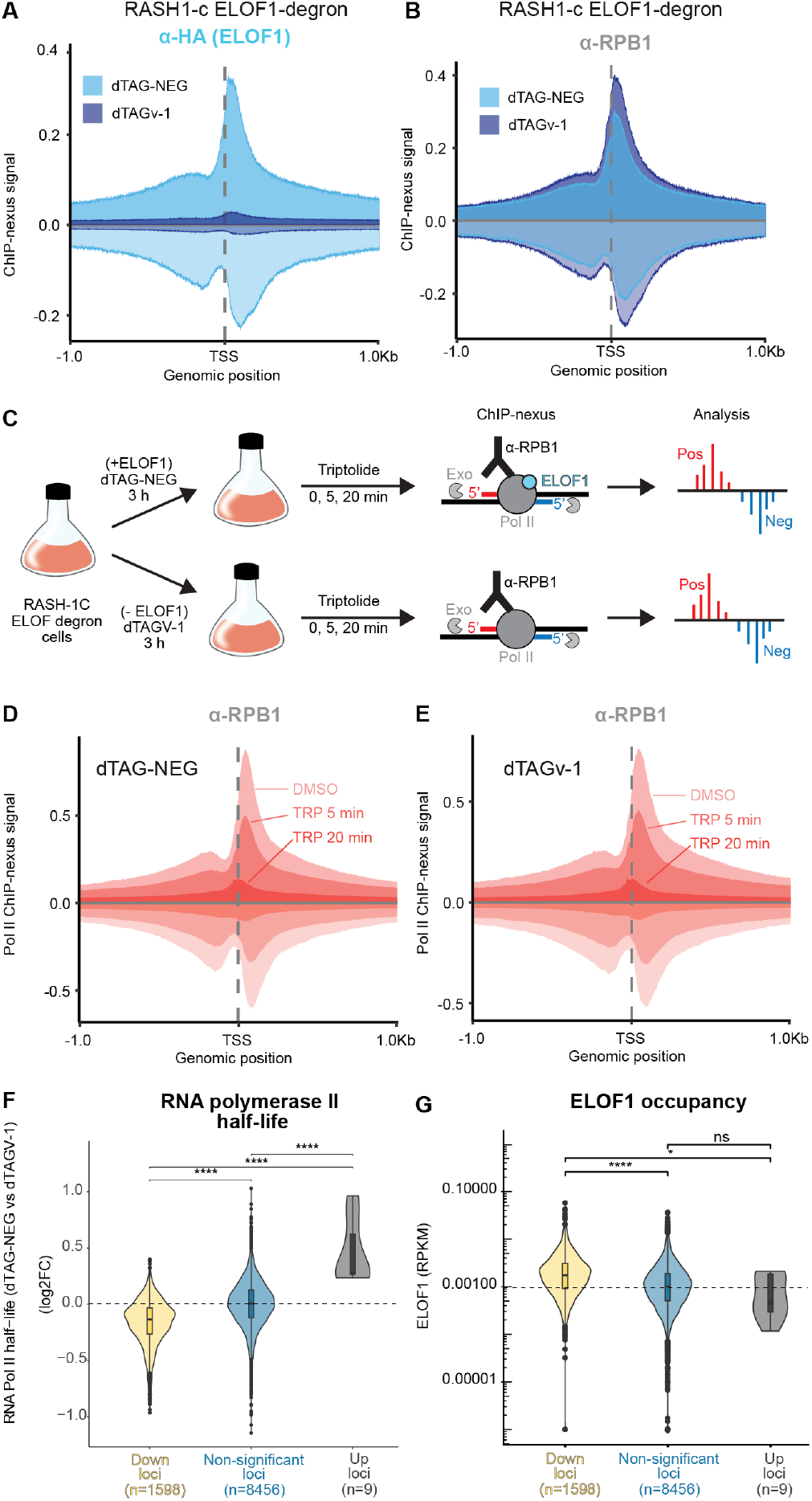
ELOF1 degradation reduces RNA polymerase II residence time in promoter-proximal regions. (**A**) Normalized ChIP-nexus metagene profiles from RASH-1C cells showing signal after 3 hours of treatment with dTAG-NEG or dTAGV-1 using anti-HA antibody to immunoprecipitate tagged ELOF1. The positive strand signal is shown above the baseline (darker shade) and the negative strand signal below the baseline (light shade). Average ChIP-nexus signal from two biological replicates is shown. (**B**) Normalized ChIP-nexus metagene profiles as in A) using anti-RPB1 N-terminal antibody. (**C**) Schematic of triptolide treatment and ChIP-nexus time course experiment in RASH-1C cells after ELOF1 degradation with dTAGV-1 or control cells treated with dTAG-NEG. (**D**) ChIP-nexus metagene profiles as in B), after 3 hours treatment with dTAG-NEG, followed by DMSO vehicle control or 5- and 20-minute treatment with Triptolide. Average signal from two biological replicates. (**E**) ChIP-nexus metagene profiles as in D) but with cells treated with dTAGV-1. (**F**) Distribution of half-life log_2_ fold-change (FC) of promoter-proximal RNA polymerase II signal in cells treated with dTAGV-1 versus cells treated with dTAG-NEG. Down, Non-significant, and Up loci were clustered based on significance of ChIP-nexus signal decay between DMSO control and 20 min of triptolide treatment in promoter-proximal regions (TSS-0.1Kb to TSS+0.5Kb). Distributions compared using Wilcoxon rank-sum test (**** p_adj_< 0.0001) (**G**) Distribution of ELOF1 ChIP-nexus signal in reads per kilobase million (RPKM) at Up, Non-significant, and Down loci from anti-HA signal shown in A). Distributions compared using Wilcoxon rank-sum test (**** p_adj_< 0.0001, * p_adj_< 0.5, ns non-significant).

Treatment with dTAGV-1 for 3 hours showed global depletion of ELOF1 from chromatin (**Fig. 1A**). In contrast, RPB1, the largest RNA polymerase II subunit, showed a mild increase in promoter-proximal occupancy upon dTAGV-1 treatment compared to dTAG-NEG treated cells (**Fig. 1B**). Such an increase is consistent with an increased initiation frequency due to shorter promoter-proximal pausing events^47,48^.

We next used ChIP-nexus to examine RPB1 and DSIF subunit SPT5 occupancy at promoter-proximal regions in ΔELOF1 human HCT116 cells and their parental line^39^. RPB1 occupancy displayed widespread, locus-specific differences (FDR < 0.05), with both gains and losses, resulting in a largely unchanged global occupancy in the promoter-proximal and gene body regions in both cell lines (**Supplementary Fig. 1C**). In contrast, SPT5, an essential protein involved in pausing and elongation, was reduced by ∼60% at pro-moter-proximal regions in ΔELOF1 cells relative to the parental cell line (**Supplementary Fig. 1D**). Because SPT5 is required for productive elongation, its loss from the promoter-proximal gene region in ΔELOF1 cells suggests that the cells have adopted compensatory adaptations that likely mask ELOF1’s direct role in transcription elongation^10,49^.We therefore did not pursue further experiments in this cell line. These results show that ELOF1 is enriched in promoter-proximal gene regions and its permanent ablation can significantly diminish the occupancy of pausing and elongation factor DSIF within promoter-proximal gene region.

### ELOF1 degradation reduces RNA polymerase II residence time in promoter-proximal regions

We next assessed if ELOF1 affected RNA polymerase II half-life in the promoter-proximal region of genes. If ELOF1 is involved in stabilizing promoter-proximal pausing, ELOF1 loss should result in a shorter paused RNA polymerase II half-life. The half-life of promoter-proximally paused RNA polymerase II can be calculated by blocking transcription initiation and assessing RNA polymerase II occupancy in promoter-proximal regions various times after transcription initiation inhibition^47,50– 52^. Here we achieved this by adding triptolide, a covalent inhibitor of TFIIH helicase XPB^53^ (**Fig. 1C**). Occupancy of RNA polymerase II at promoter-proximal regions decays over time due to its release into processive elongation or its premature termination. RPB1 occupancy was measured by ChIP-nexus in ELOF1-degron cells (**Fig. 1C**). Cells were treated for 3 hours with dTAGV-1 or dTAG-NEG to ensure ELOF1 depletion prior to adding triptolide. Samples were taken at 0, 5, and 20 minutes after triptolide addition (**Fig. 1C**).

As expected, triptolide addition resulted in the time-dependent loss of RPB1 signal within the promoter-proximal regions of genes with no indication of RPB1 degradation (**Fig. 1D, E, Supplementary Fig. 1E**). Normalized ChIP-nexus counts were fit to a single exponential decay to determine RPB1 half-lives at all promoter-proximal regions (**Supplementary Fig. 1F**). The distribution of half-lifes, with a median half-life of ∼7 min, resembled previously reported values estimated in *Drosophila*, mouse, and human cultured cells, (**Supplementary Fig. 1G**)^47,50,51^. We then compared half-life values in dTAGV-1 and dTAG-NEG treated cells (log_2_ Fold Change (FC)). Loci showing a significant reduction in RPB1 occupancy (“down-loci”; log_2_FC < 0 and p_adj_ < 0.05) displayed a median half-life log_2_FC of -0.14 indicating faster RNA polymerase II turnover at promoter-proximal positions in the absence of ELOF1 (**Fig. 1F**). No change was observed at loci with unchanged or increased RPB1 occupancy (**Fig. 1F**). Down-loci also showed ∼1.7 fold higher ELOF1 occupancy in dTAG-NEG treated cells than the other two groups, indicating a strong link between pause stability and ELOF1 occupancy (**Fig. 1F, G**). Finally, Gene Ontology (GO) analysis of the down-loci group produced no significant enrichment of any gene class or overlap with genes targeted by AID during somatic hypermutation, a key function of RASH-1C cells. Together these results indicate that ELOF1 helps to maintain paused RNA polymerase II and that ELOF1 is a general rather than a gene or cell-type specific transcription elongation factor.

### ELOF1 strongly enhances promoter-proximal pausing induced by DSIF and NELF

To further probe the role of ELOF1 in promoter-proximal pausing by RNA polymerase II, we performed Gene-specific Analysis of Transcriptional Output (GATO-seq). GATO-seq is a cell-free genomics assay that allows for the time-resolved and modular assessment of RNA synthesis by RNA polymerases on a library of DNA sequences by mapping nascent RNA 3’ ends using direct RNA Oxford Nanopore sequencing^54^ (**Fig. 2A**).

**Figure 2.**
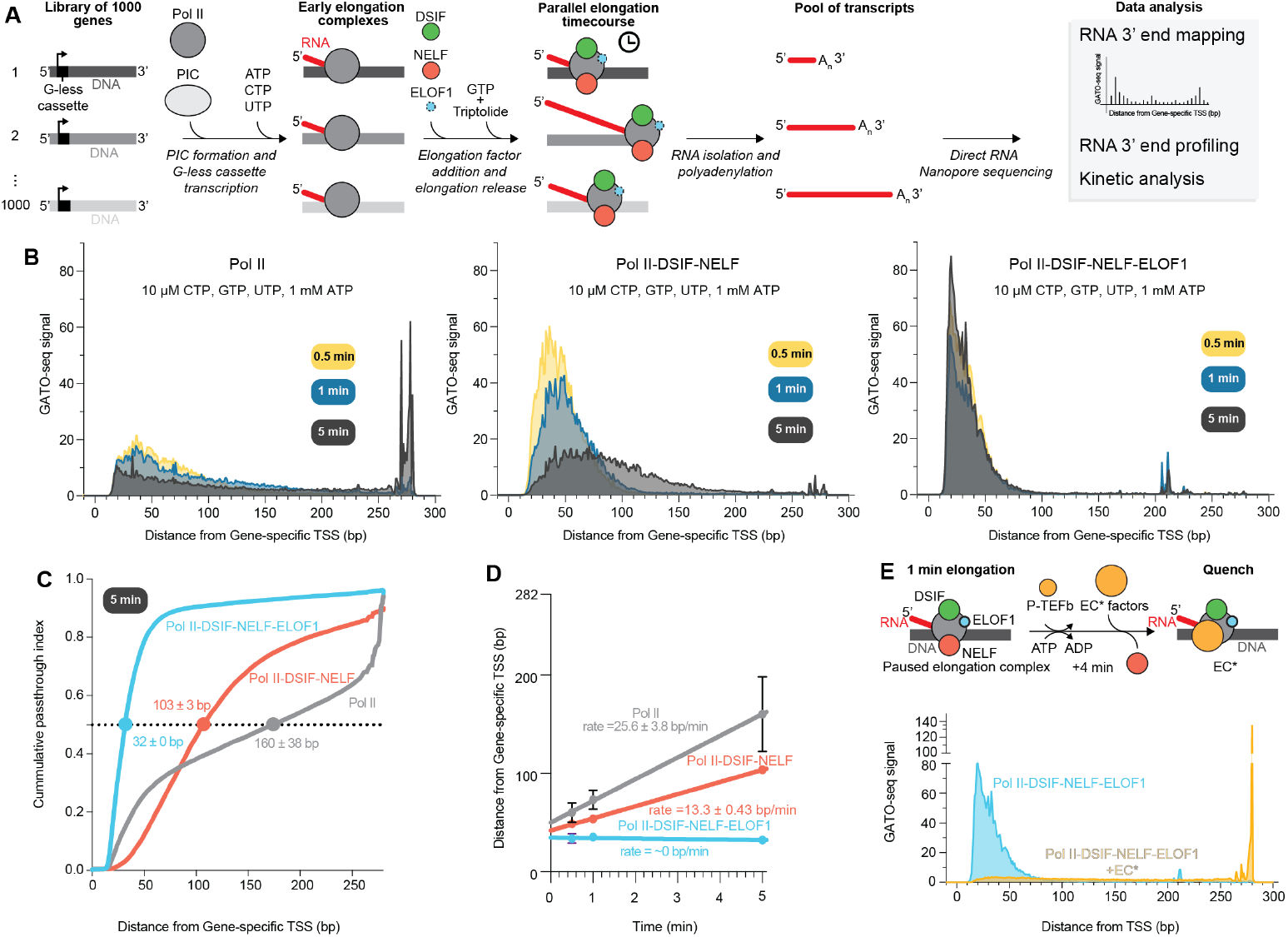
ELOF1 strongly enhances promoter-proximal pausing. (**A**) Schematic of GATO-seq experimental design to test addition of elongation factors DSIF, NELF and ELOF1 on transcription elongation activity. (**B**) Metagene profiles of GATO-seq signal at three different time points from two biological replicates employing 10 µM CTP, GTP, UTP and 1 mM ATP. Left, RNA polymerase II without additional factors from^54^, middle with 3-fold molar excess of DSIF and NELF from^54^ and right with 3-fold molar excess of DSIF and NELF and 10-fold molar excess of ELOF1. Average signal from two biological replicates. (**C**) Comparison of passthrough index cumulative distributions at 5-minute time point from two biological replicates with 10 µM CTP, GTP, UTP and 1 mM ATP and in the absence of elongation factors. (Pol II)^54^, with DSIF and NELF^54^, and DSIF, NELF and ELOF1. Circles indicate cumulative distribution midpoint (y=0.5). (**D**) Cumulative distribution midpoint as a function of time using 10 µM CTP, GTP, UTP and 1 mM ATP. Rates were estimated from a linear fit (Pol II = 25.6 ± 3.8 bp/min, Pol II-DSIF-NELF = 13.3 ± 0.43 bp/min and Pol II-DSIF-NELF-ELOF1 = ∼ 0 bp/min). Error bars represent the standard deviation between two biological replicates. (**E**) Top: schematic of pause release from the RNA polymerase II-DSIF-NELF-ELOF1 complex to the activated EC* complex (P-TEFb and PAF1c (with RTF1), SPT6 and TFIIS). Bottom: Metagene profiles of GATO-seq signal. Elongation proceeded for 1 minute with DSIF, NELF, and ELOF1, then for 4 additional minutes ± EC* factors (5 minutes total). Average signal from two biological replicates.

We compared the effect of ELOF1, DSIF, NELF, or combinations of these factors on RNA polymerase II promoter-proximal pausing using a library of 1000 human promoter-proximal DNA sequences encoding the first 262 bases of each gene. Each gene body sequence was appended to the Adenovirus Major Late (AdML) promoter and a 9 bp G-less cassette to facilitate the coordinated formation of early transcription elongation complexes (**Supplementary Fig. 2A**). RNA polymerase II, transcription initiation factors, and ATP, CTP, and UTP were first incubated with the library to permit transcription initiation and the synthesis of the G-less cassette. GTP and triptolide were next added to allow for time-resolved, single turnover transcription elongation events (**Fig. 2A**). Initial experiments were performed under a low nucleotide regime to enhance detection of promoter-proximal pausing (10 µM CTP, GTP, UTP, and 1 mM ATP for TFIIH helicase and kinase activities). Transcription elongation proceeded for 30 s, 60 s, and 5 minutes prior to quenching the reactions. Metagene plots of the normalized RNA 3’ end counts (GATO-seq signal) from RNA polymerase II samples containing no additional elongation factors showed progression of RNA polymerase II across the gene bodies over time (**Fig. 2B**). Addition of DSIF and NELF slowed the traversal of RNA polymerase II across the gene body whereas ELOF1 addition closely mirrored RNA polymerase II samples without additional elongation factors (**Fig. 2B, Supplementary Fig. 2B, C**). In contrast, a single peak centered around ∼20 bp was observed at all time points when ELOF1 was combined with DSIF and NELF (**Fig. 2B**). We next defined the relative position of RNA polymerase II at each time point using the passthrough index, which accounts for the upstream signal relative to the total signal at each base position^54^ (**Supplementary Fig. 2D**). The cumulative passthrough index distribution reports at the midpoint (y=0.5) the average position of RNA polymerase II at a given time point. (**Supplementary Fig. 2E**). After 5 minutes of transcription, RNA polymerase II reached an average position of 160 ± 38 bp, which decreased to 103 ± 3 bp when DSIF and NELF were included^54^ (**Fig. 2C**). This position was shifted to 32 ± 0 bp when ELOF1 was included with DSIF-NELF (**Fig. 2C**). The relative velocity of RNA polymerase II was estimated using the change in passthrough index midpoint over time. RNA polymerase II without additional elongation factors had a relative rate of 25.6 ± 3.8 bp/min^54^. DSIF and NELF reduced this rate to 13.3 ± 0.43 bp/min, whereas no translocation was detected (rate ∼ 0 bp/min) when DSIF-NELF-ELOF1 were included (**Fig. 2D**).

These results suggested that ELOF1 either enhanced pausing induced by DSIF and NELF or supported premature termination. To define if RNA polymerase II was still engaged with nucleic acids in the presence of DSIF-NELF-ELOF1, we tested whether RNA polymerase II could be released from pausing into productive elongation by adding P-TEFb and elongation factors PAF1c (here with RTF1), SPT6, and TFIIS to form the activated elongation complex (EC*)^15,16,55^. We allowed RNA polymerase II to elongate for 1 minute in the presence of DSIF-NELF-ELOF1, followed by addition of EC* factors with 1 mM ATP. Transcription elongation proceeded for an additional 4 minutes before quenching (**Fig. 2E**). Addition of EC* elongation factors resulted in a shift of the passthrough index midpoint to 183 ± 16 bp (**Fig. 2E, Supplementary Fig. 2E**). These results showed that DSIF-NELF-ELOF1 strongly slowed RNA polymerase II and did not lead to substantial premature termination.

### DSIF-NELF-ELOF1 potently induce RNA polymerase II pausing under physiological conditions

In reconstituted systems, DSIF and NELF have a limited role in reducing RNA polymerase II synthesis rates at physiological nucleotide concentrations^30,69–71^. W^66^e thus assessed whether the inclusion of ELOF1 altered this behavior. GATO-seq experiments were performed with 1 mM of each NTP. Under these conditions, the average position of RNA polymerase II was at 172 ± 11 bp without additional elongation factors after 30 s of transcription elongation (**Fig. 3A, B, Supplementary Fig. 2F, G**). In the presence of DSIF-NELF this value corresponded to 114 ± 2 bp (**Fig. 3A, B, Supplementary Fig. 2F, G**). Strikingly, reactions containing DSIF-NELF-ELOF1 resulted in an average position of 55 ± 1 bp (**Fig. 3A, B, Supplementary Fig. 2F, G**). The relative velocity of RNA polymerase II in these different conditions was estimated using the average position of RNA polymerase II at different time points (**Fig. 3B, Supplementary Fig. 2G**). RNA polymerase II without additional elongation factors or with DSIF-NELF had relative elongation rates of ∼112 bp/min and ∼52 bp/min, respectively (**Fig. 3C**). Given that most RNA polymerase II reached the end of templates after 30 s, we are likely underestimating the elongation rate in both conditions. Addition of DSIF-NELF-ELOF1 significantly slowed down RNA polymerase II progression (29.08 ± 0.43 bp/min) (**Fig. 3C**). Together, these results show that ELOF1 acts synergistically with DSIF and NELF to decrease the rate of transcription elongation and that its inclusion with DSIF-NELF reduced elongation rates at physiological nucleotide concentrations.

**Figure 3.**
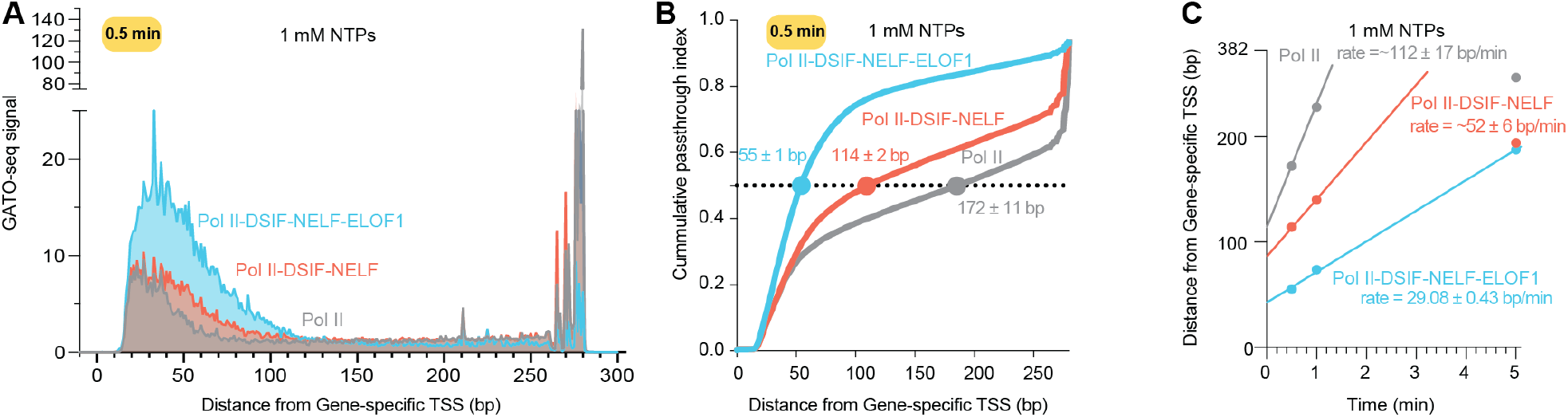
ELOF1 stabilizes promoter-proximal pausing at physiological NTP concentrations. (**A**) Metagene profiles of GATO-seq signal at 30 seconds from two biological replicates employing 1 mM NTPs. (**B**) Comparison of passthrough index cumulative distributions at 30 seconds from two biological replicates with 1 mM NTPs and in the absence of elongation factors (Pol II^54^), with DSIF-NELF, and DSIF-NELF-ELOF1. Circles indicate cumulative distribution midpoint (y=0.5). (**C**) Cumulative distribution midpoint as a function of time using 10 µM CTP, GTP, UTP and 1 mM ATP. Rates were estimated from a linear fit (Pol II = 112 ± 17 bp/min, Pol II -DSIF-NELF = 52 ± 6 bp/min and Pol II-DSIF-NELF-ELOF1 = 29 ± 0.4 bp/min). Error bars represent the standard deviation between two biological replicates. Some error bars are not visible due to small values. Dotted line indicates the end of the template.

### ELOF1 reduces pause escape rate

Pausing is a transient state, with only a fraction of RNA polymerase molecules entering the paused state at a given pause site. This can be defined as the pause efficiency (E)^56,57^. If RNA polymerase II does pause, the escape into elongation can be measured as the pause escape rate (t_1/2_)^56^. From GATO-seq experiments, the pause efficiency and the pause escape rate can be estimated at pause sites that show decay in signal over time^54^. We thus assessed whether addition of DSIF-NELF with or without ELOF1 altered the pause position, pause efficiency or escape rate compared to RNA polymerase II in the absence of these factors. Pause sites correspond to positions where GATO-seq signal is ≥4 standard deviations above the gene average in at least two consecutive time points^54^. Using these criteria and our low nucleotide data, a similar number of pause sites were detected for DSIF-NELF (2718) and DSIF-NELF-ELOF1 (2741 sites) samples whereas 1702 sites were detected in RNA polymerase II without additional elongation factors (**Fig. 4A**). The number of detected pause sites was greatly diminished in reactions containing 1 mM of NTPs (516 sites RNA polymerase II without additional elongation factors, 897 sites DSIF-NELF, 1588 sites DSIF-NELF-ELOF1) (**Fig. 4A**).

**Figure 4.**
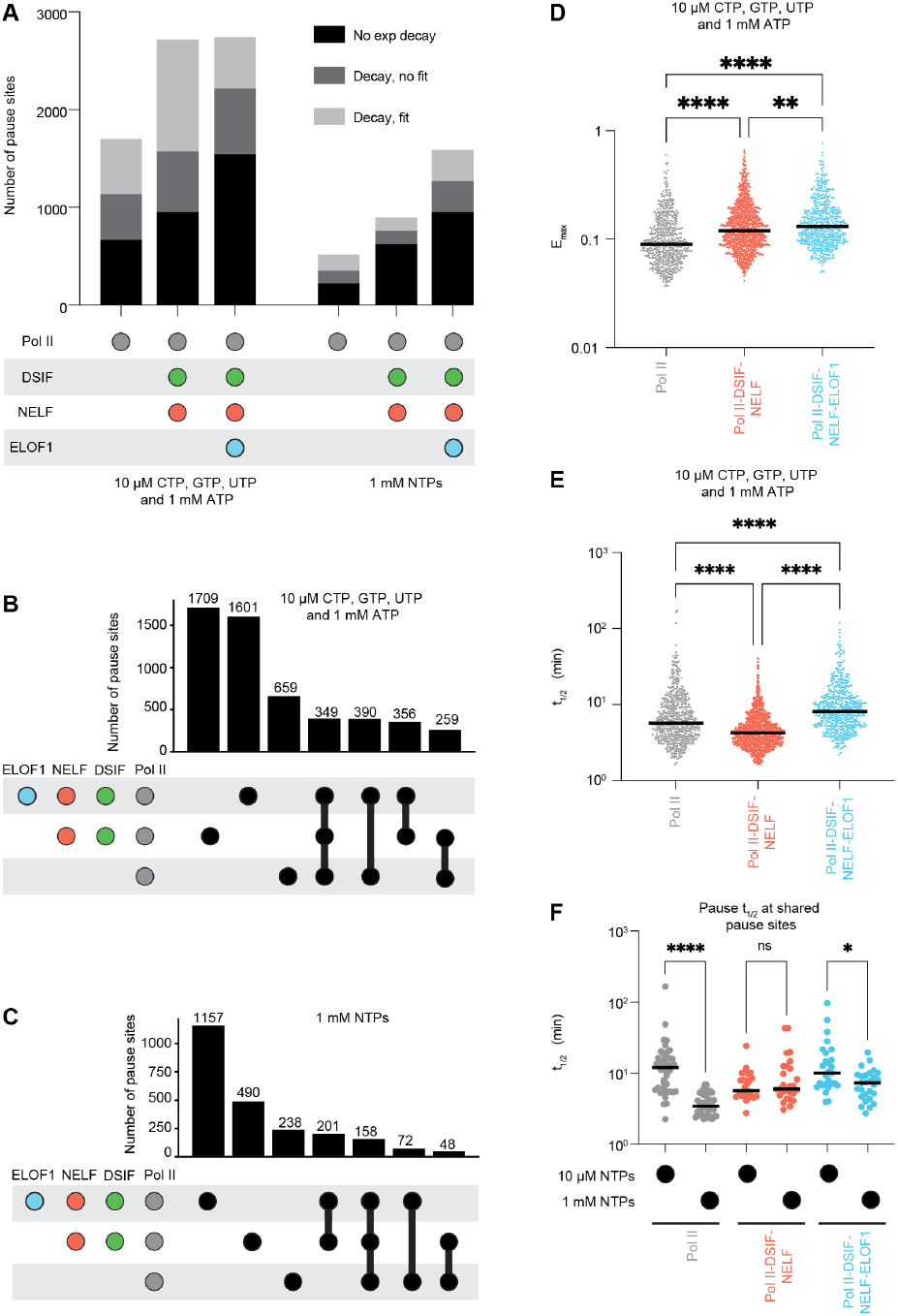
ELOF1 reduces pause escape rate. (**A**) Sequence logo derived from TFIIS insensitive sites (n=43 sequences). (**B**) Super pause consensus sequence scaffold used for RNA extension assays (**C**) RNA primer extension time course experiments in the absence or presence of TFIIS and 10 µM CTP, GTP and UTP. RNA polymerase II:TFIIS used at a molar ratio of 1:0.2. All experiments were performed at least three times on super pause consensus scaffold (**D**) RNA extension assay with 1 mM CTP, GTP and UTP on the super pause scaffold and the super pause scaffold with a +1G to U substitution. (**E**) RNA extension assay as in (C) with super pause scaffold caring a +1G to U substitution (**F**) RNA extension assay as in (C) caring a -2C to U substitution.

Next, we compared the position of pause sites across all conditions. Only a fraction of pause sites overlapped. We first analyzed pause positions that were retained in all factor combinations and nucleotide concentrations. A total of 394 and 158 pause sites were detected in all conditions at low and high NTP concentrations, respectively (**Fig. 4B, C**).

We next measured the maximal pause efficiency (E_max_), which corresponds to the y intercept. The median E_max_ value of RNA polymerase without elongation factors was 0.089, and it increased to 0.11 when DSIF-NELF were present (**Fig. 4D**). The median E_max_ further increased to 0.13 when DSIF-NELF-ELOF1 were present (**Fig. 4D**). DSIFNELF-(ELOF1) thus appear to enhance the proportion of RNA polymerase II molecules that enter the paused state. Finally, we compared the pause escape half-life (t_1/2_). At low NTP concentrations, addition of DSIF-NELF decreased the median t_1/2_ to 4.27 min whereas RNA polymerase II in the absence of elongation factors had a median t_1/2_ of 5.71 min^54^ (**Fig. 4E**). When ELOF1 was included with DSIF-NELF the median half-life increased to 8.13 min, indicating that addition of ELOF1 to RNA polymerase II-DSIF-NELF complexes decreased the pause escape rate (**Fig. 4E**). To address the sensitivity of t_1/2_ to NTP concentration, we compared pause sites overlapping between low and high NTP conditions. In the absence of elongation factors, the t_1/2_ was reduced from 12 min to 3.4 min from the low and high NTP concentration samples, respectively (**Fig. 4F**). The t_1/2_ displayed a negligible increase from 5.6 to 6 minutes when DSIF-NELF were present whereas addition of DSIF-NELF-ELOF1 resulted in a change from 10 min to 7.2 minutes, between low and high nucleotide samples, respectively. These results suggest that without DSIF-NELF and/or ELOF1, RNA polymerase II release from pausing is highly correlated with NTP concentration (**Fig. 4F**) and DSIF-NELF- and/or ELOF1 buffer this effect to differing extents. Together, ELOF1 in combination with DSIF-NELF decreases the translocation rate of RNA polymerase II, enhances entry into a paused state (E_max_), increases the number of detected pause sites, and decreases the pause escape rate (t_1/2_).

### Cryo-EM structure of RNA polymerase II-DSIF-NELF-ELOF1 complex

To understand how ELOF1 associates with the RNA polymerase II-DSIF-NELF complex, we determined the cryo-EM structure of the complex on a nucleic acid scaffold that induces strong pausing behavior^54^. This perfectly complementary scaffold comprises 13 bp of upstream DNA, a 9 bp DNA-RNA hybrid positioned 3 bp upstream of the pause site, 33 bp of downstream DNA and 6 nt of exiting RNA (**Supplementary Fig. 3A**). RNA polymerase II was assembled on the nucleic acid scaffold with DSIF, NELF, and ELOF1 and permitted to transcribe for 10 minutes prior to cryo-EM sample preparation (**Supplementary Fig. 3B, C and D**). Using actively transcribing complexes allowed for the capture of various conformational states of DSIF and NELF and for RNA polymerase II to sample distinct translocation registers^58^. This is important because RNA polymerase pausing is known to frequently involve offline active site states^54,58,59,66,67^. After transcription, predominant RNA species included the starting RNA primer and a +3 extended or paused species (**Supplementary Fig. 3D**).

Cryo-EM data was collected, and a total of 8,251,315 particles were extracted (**Supplementary Fig. 3E, F**).

Subsequent 2D and 3D classification produced distinct compositional and conformational states (**Supplementary Fig. 4 and 5**). A map containing RNA polymerase II, DSIF, NELF in the paused conformation, and ELOF1 was obtained from 84,105 particles with a resolution ranging from 2.7-8 Å (gold-standard 0.143 Fourier Shell Correlation (FSC) cutoff) (**Fig. 5A-C, Supplementary Fig. 5A, B, Table 1**). We additionally classified a subset of 134,424 particles containing RNA polymerase II-DSIF-NELF-ELOF1 with NELF in the poised conformation^58,60^ (**Supplementary Fig. 4**). The poised NELF conformation showed more conformational heterogeneity and was thus not used for modelling. Importantly, both maps show the same features for RNA polymerase II, DSIF, and ELOF1. Additional maps were obtained for

**Figure 5.**
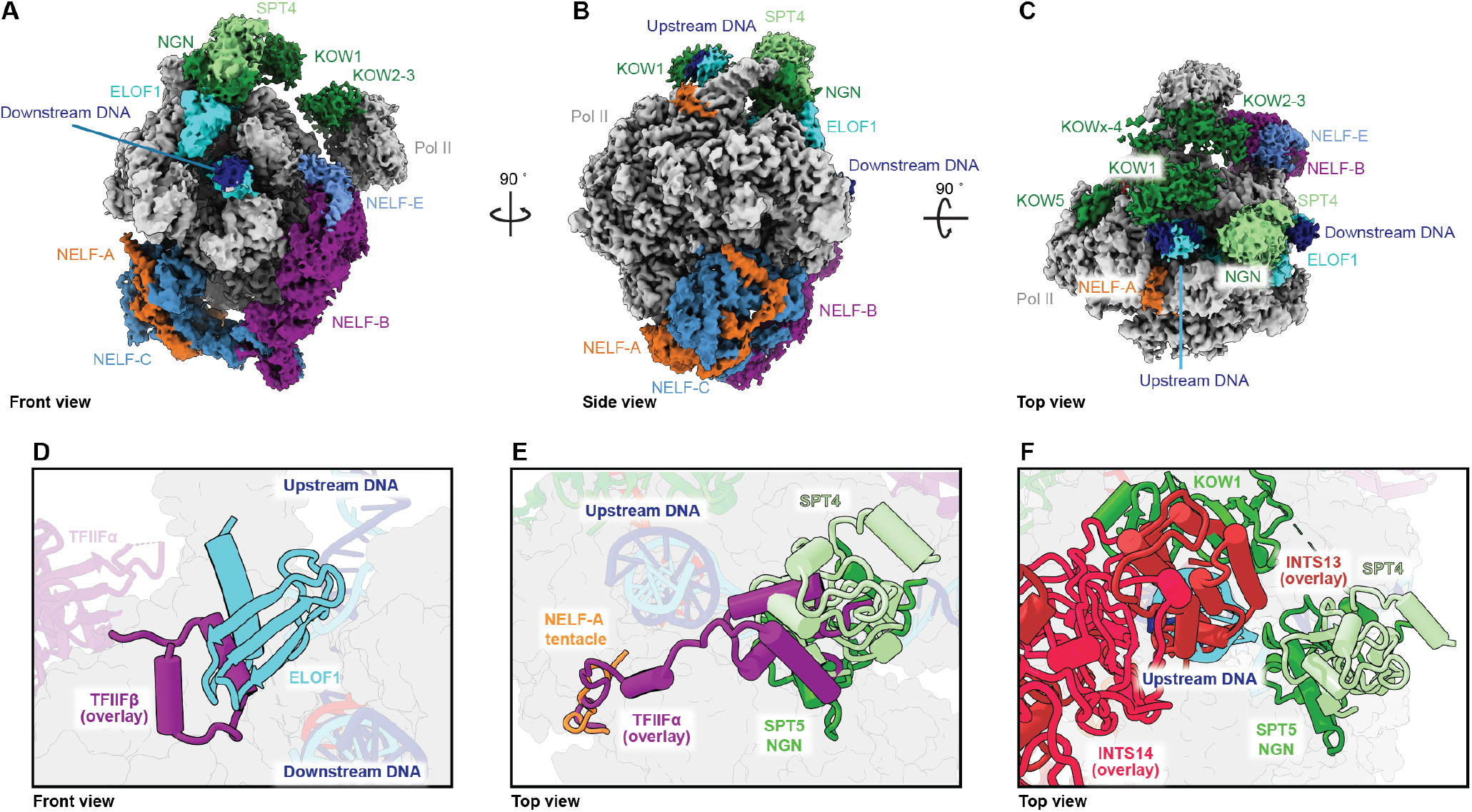
Cryo-EM structure of RNA polymerase II-DSIF-NELF-ELOF1 paused elongation complex. (**A-C**) RNA polymerase II-DSIF-NELF-ELOF1 cryo-EM composite-map. RNA polymerase II is colored silver, template DNA blue, non-template DNA cyan, and RNA red. NELF subunits are colored as: NELF-A orange, NELF-C marine, NELF-B purple, and NELF-E light blue. SPT4 is colored light green and SPT5 dark green. ELOF1 is shown in light blue. (A) Front, (B) side, and (C) top views are shown. (**D**) Front view, overlay of preinitiation complex TFIIF (PDB 7NVU) and RNA polymerase II-DSIF-NELF-ELOF1 structural model. TFIIF-shown in orchid, and TFIIF-in purple, other colors as in panels A-C. Dotted circle indicates a steric clash between ELOF1 and TFIIF. (**E**) Top view, overlay as in B). Dotted circles indicate steric clashes between TFIIF and NELF-A tentacle, SPT4, and SPT5 NGN domain. (**F**) Side view, overlay of Integrator pretermination complex (PDB 8RBX) and RNA polymerase II-DSIF-NELF-ELOF1 structural model. INTS13 and INTS14 subunits are shown in brown and crimson colors, respectively. Other colors as in panels A-C. Dotted circle indicates clashes between INTS13 and upstream DNA and SPT5 KOW1 domain.

**Table 1:** Cryo-EM data collection, refinement, and validation statistics.

|  | RNA polymerase II-DSIF-NELF-ELOF1 elongation complex (EMD-XXXX) (PDB: XXXX) |
| --- | --- |
| **Data collection and processing** |  |
| Magnification | 105,000 |
| Voltage (kV) | 300 |
| Electron exposure (e-/Å<sup>2</sup>) | 50.1 |
| Defocus range (µm) | 0.25-2.25 |
| Pixel size (Å) | 0.822 |
| Symmetry imposed | C1 |
| Initial particle images (no.) | 8,251,315 |
| Final particle images (no.) | 84,105 |
| Map resolution (Å) | 2.7 |
| FSC threshold | 0.143 |
| Map resolution range (Å) | 2.7-8 |
| **Refinement** |  |
| Initial models used (PDB code) | 9BZ0, 8UHD, and 3H7H |
| Map sharpening B factor (Å<sup>2</sup>) | -15 |
| **Model composition** |  |
| Non-hydrogen atoms | 94008 |
| Protein residues | 5756 |
| Nucleotides | 87 |
| Ligands | 11 |
| **B factors (Å<sup>2</sup>)** |  |
| Protein | 111.63 |
| Nucleotide | 228.03 |
| Ligand | 183.23 |
| **R.M.S. deviations** |  |
| Bond lengths (Å) | 0.003 |
| Bond angles (°) | 0.517 |
| **Validation** |  |
| MolProbity score | 1.5 |
| Clashscore | 4.87 |
| Poor rotamers (%) | 0.71 |
| **Ramachandran plot** |  |
| Favored (%) | 96.32 |
| Allowed (%) | 3.68 |
| Disallowed (%) | 0 |

RNA polymerase II without additional factors, RNA polymerase II with either ELOF1 or DSIF, RNA polymerase II-DSIF-NELF, and RNA polymerase II-DSIF-ELOF1 (**Supplementary Fig. 4 and 5**).

High resolution features allowed for unambiguous assignment of ELOF1 in our cryo-EM maps. The ELOF1 binding mode on RNA polymerase II largely resembled previously determined structures of activated elongation complexes (RMSD 1.92-2.38 Å) and is shifted ∼5 Å towards the downstream DNA relative to its location in TCR-NER structures (RMSD 4.02 Å) ^41–43,61–64^ (**Supplementary Fig. 6A, B**). ELOF1 comprises three antiparallel beta strands that contact the RPB1 clamp and a cysteine coordinated zinc finger that is preceded by a C-terminal alpha helix that anchors ELOF1 to the RPB2 lobe. This arrangement forms a downstream DNA entry tunnel that has been proposed to support processive elongation in activated elongation complexes^41–43,61,65^. Our maps lacked density for the first 15 residues of ELOF1 as previously observed ^41,43,61,62^. The positions of DSIF and NELF on RNA polymerase II are nearly identical to previously determined structures without ELOF1^58–60^ (**Fig. 5A-C**). No direct interactions between ELOF1 and NELF were observed.

The SPT4-SPT5 NusG N-terminal (NGN) domain is dynamic in cryo-EM structures lacking ELOF1, as also observed here, and has required focused classifications and refinements to visualize it^58–60^. Strikingly, maps containing DSIF and ELOF1 with or without NELF showed strong density for SPT4-SPT5 NGN and required no additional classification (**Fig. 5A-C, Supplementary Fig. 5B and 6C**). As previously observed, ELOF1 directly contacts the SPT5 NGN (**Fig. 5A-C, Supplementary Fig. 6C**)^42,43,62^. This highlights that the ELOF1-SPT5 NGN domain interface is retained in both processive and paused transcription elongation complexes and that ELOF1 stabilizes the SPT4-SPT5 NGN. Overlaying structures of transcription initiation complexes onto our RNA polymerase II-DSIF-NELF-ELOF1 structure shows that the position of NELF-SPT4-SPT5-NGN-ELOF1 sterically clashes with the positions of initiation factors TFIIF and TFIIE on RNA polymerase II (**Fig. 5D, E, Supplementary Fig. 6D, E**). We next overlaid Integrator from a pretermination complex onto our structure. Interestingly, the Integrator tail module subunit INTS13 and the upstream DNA clamp formed by SPT4-SPT5-NGN-KOW1 are structurally incompatible^24^(**Fig. 5F**).

Finally, we assessed the nucleic acid register within the RNA polymerase II active site. Pausing is associated with offline active site states including backtracking and tilted/half-translocated states^58,59,66,67^. The RNA polymerase II-DSIF-NELF-ELOF1 complex was predominantly in a posttranslocated active site conformation (**Supplementary Fig. 6F**). The nucleic acid sequence used in our cryo-EM studies can support an offline active site state known as ‘sidetracking’^54^. No density for side-tracked or backtracked RNA was detected in maps containing RNA polymerase II-DSIF-NELF with or without ELOF1. In contrast, the RNA polymerase II-DSIF and RNA polymerase II-DSIF -ELOF1 maps showed a subset of particles with a sidetracked RNA 3’ end (**Supplementary Fig. 4 and 6G**). There are two explanations for this observation (1) RNA polymerase II-DSIF-NELF-(ELOF1) did not reach the pause site or (2) the association of DSIF-NELF with RNA polymerase II prevents sidetracking. The heterogeneity of the actively transcribing complex used for cryo-EM sample preparation prevented unambiguous assignment of the nucleic acid sequence within the active site, and thus we cannot presently clarify which of these models is correct. Overall, cryo-EM structures of RNA polymerase II-DSIF-NELF-ELOF1 show that ELOF1 does not alter the conformation of the paused complex and that ELOF1 stabilizes the SPT5 NGN as previously observed in other transcription elongation complexes.

### ELOF1 stabilizes DSIF and NELF on RNA polymerase II

TFIIF is an initiation factor that can associate with elongating RNA polymerase II, enhance its processivity, and counteract the negative effects of DSIF-NELF^30,31^. TFIIF shares overlapping binding surfaces with DSIF-NELF on RNA polymerase II. Further, functional assays have shown competition between TFIIF and DSIF-NELF^31^. Because ELOF1 partially shares a binding surface with TFIIF and enhances DSIF-NELF dependent pausing, we hypothesized that ELOF1 is the “TFIIF resistance factor” that supports DSIF-NELF dependent pausing of RNA polymerase II in the presence of TFIIF. If this hypothesis is true, the DSIF-NELF-ELOF1 complex should be resistant to TFIIF addition.

To test this hypothesis, we performed RNA extension assays with RNA polymerase II in the presence of different protein factor combinations. We utilized the bacterial elemental pause sequence where mammalian RNA polymerase II displays pausing, backtracking, and processive elongation behaviors. Specifically, RNA polymerase II pauses after adding two nucleotides to the RNA primer, can back-track after incorporating three nucleotides, and achieves maximum extension at seven nucleotides in the presence of GTP and CTP^58,59,68^ (**Supplementary Fig. 6H**).

As previously observed, DSIF-NELF slowed RNA polymerase II elongation activity whereas TFIIF greatly enhanced this activity^58,59^ (**Fig. 6A, B**). TFIIF addition to DSIF-NELF containing reactions fully counteracted the negative effects of DSIF-NELF^30,31^(**Fig. 6A, B**). Having confirmed previous reports, we next included ELOF1 in our assays. ELOF1 addition to DSIF-NELF containing reactions exhibited similar behaviors as reactions containing only DSIF-NELF (**Fig. 6A, B**). In contrast, DSIF-NELF-ELOF1 reactions showed sustained pausing behavior in the presence of TFIIF (**Fig. 6A, B**). This TFIIF suppression behavior was only achieved when DSIF, NELF and ELOF1 were all present (**Supplementary Fig. 6I, J**). These functional experiments show that ELOF1 operates as a resistance factor to TFIIF when combined with DSIF-NELF.

**Figure 6.**
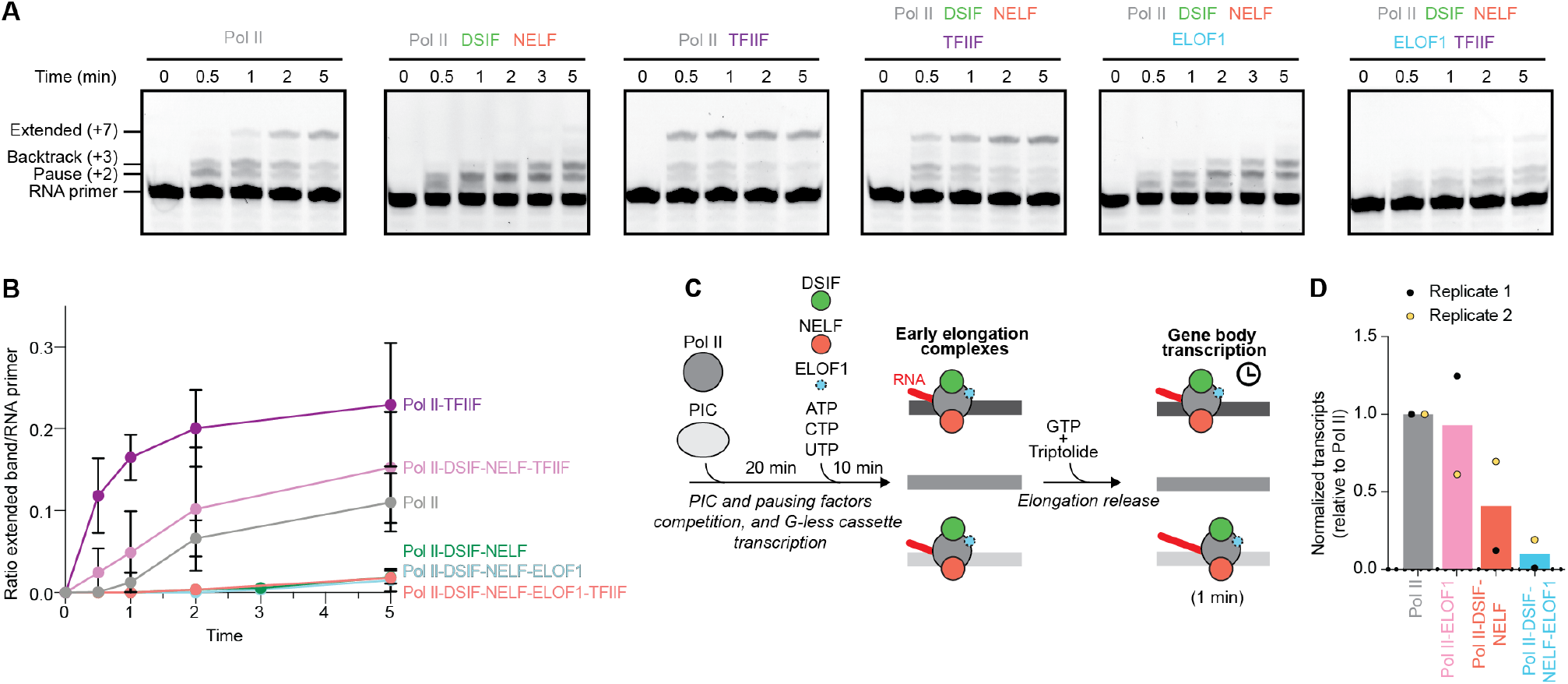
ELOF1 confers resistance to TFIIF in the presence of DSIF and NELF. (**A**) RNA primer extension time course experiments with 10 µM CTP and GTP and in the absence or presence of indicated elongation factors used in the following ratios relative to RNA polymerase II: TFIIF 1:5, DSIF 1:3, NELF 1:3, ELOF1 1:10. All experiments were performed at least three times, and a representative gel is shown. (**B**) Intensity ratio of extended (+7) band relative to the RNA primer quantified from A). (**C**) Schematic GATO-seq experiment to monitor competition between elongation and initiation factors. (**D**) Number of normalized transcripts at indicated experimental conditions from GATO-seq competition assay. Values from two biological replicates are shown. Bar indicates average of replicates.

Having established that DSIF-NELF-ELOF1 block TFIIF activity during transcription elongation, we next sought to understand if DSIF-NELF-ELOF1 affects TFIIF’s function in the preinitiation complex (PIC). To test this idea, we performed GATO-seq and measured total RNA production after 1 minute of transcription elongation as a proxy for transcription initiation efficiency. To allow for competition between initiation and elongation factors we modified the order of addition of factors. DSIF, NELF and ELOF1 were included with NTPs and transcription initiation factors during transcription initiation and transcription of the G-less cassette (**Fig. 6C**). For each sample, the total amount of RNA produced was normalized to the total amount of RNA produced by RNA polymerase II in the absence of elongation factors termed “transcriptional output”. Addition of only ELOF1 had a minor effect on transcriptional output (**Fig. 6D**). DSIF and NELF led to a ∼2-fold decrease in transcriptional output, consistent with competition with initiation factors (**Fig. 6D**). When DSIF, NELF and ELOF1 were included together, transcriptional output decreased by at least 10-fold, consistent with ELOF1 enhancing DSIF-NELF function by displacing initiation factors (**Fig. 6D**). In summary, ELOF1 in combination with DSIF-NELF was sufficient to counteract the positive effects of TFIIF on RNA polymerase II activity.

## Discussion

Despite over two decades of efforts, reconstituted systems have failed to recapitulate promoter-proximal pausing under physiological conditions. Early work found that DSIF and NELF could induce pausing, but this often required limiting nucleotide concentrations^30,69–71^. This pausing was also readily reversed by addition of TFIIF^31^. This led to the hypothesis of a missing “TFIIF resistance factor”, but the identity of this factor has remained elusive^30,31^. Here, using cellular, biochemical, and structural approaches, we have identified ELOF1 as the TFIIF resistance factor. ELOF1 is enriched in the promoter-proximal region of genes, and its rapid degradation leads to reduced pause duration in cells. In reconstituted assays, ELOF1 synergizes with DSIF and NELF to induce stable promoter-proximal pausing under physiological nucleotide conditions. Cryo-EM structures reveal that DSIF-NELF-ELOF1 occupy binding surfaces on RNA polymerase II that overlap with those of TFIIF and TFIIE and Integrator. Finally, functional assays show that ELOF1 makes DSIF-NELF complexes resistant to TFIIF addition. Together, these results answer a long-standing question in the field of how promoter-proximal pausing is stably induced under physiological conditions.

Our results provide a model for how RNA polymerase II transitions from initiation into pausing (Fig. 7). Biochemical and structural studies have shown that initiation factors TFIIE and TFIIF remain bound to RNA polymerase II after transcription initiation and promoter escape^72,73^. TFIIF may be removed by PAF1c during EC* formation^73^; however, EC* formation occurs after pause release, meaning TFIIF would remain associated with RNA polymerase II throughout the pausing window and counteract DSIF-NELF. Here, we show that ELOF1 with DSIF-NELF are sufficient to counteract TFIIF. How DSIF-NELF-ELOF1 are prevented from binding RNA polymerase II during PIC assembly remains unclear. Consistent with direct competition, we observe that addition of DSIF-NELF-ELOF1 to initiation reactions nearly abolishes RNA production, highlighting that spatial or temporal separation of initiation and pausing factor binding may be critical^74–76^. Importantly, our experiments used the minimal general transcription factors (TFIIA, B, E, F, H and TBP) required to assemble the PIC, leaving open the possibility that additional components, such as the TFIID complex or Mediator, may modulate or prevent this competition in cells.

**Figure 7.**
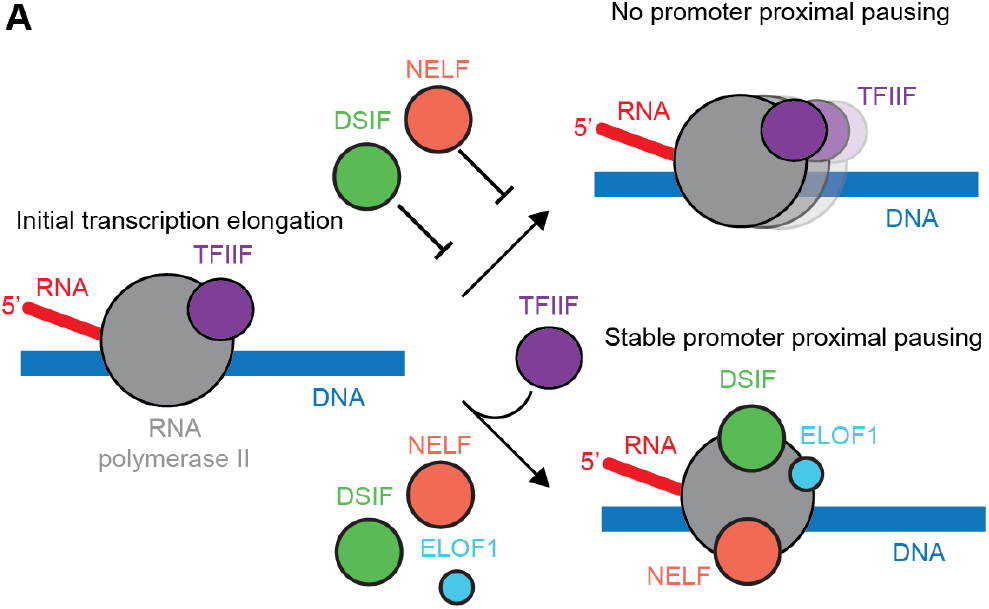
Model of action of ELOF1 in promoter-proximal pausing. During early elongation, TFIIF remains bound to RNA polymerase II. In the absence of ELOF1, the elongation factors DSIF and NELF are unable to establish promoter-proximal pausing, presumably because TFIIF hinders their binding or because they weakly associate with the RNA polymerase II-TFIIF complex. In the presence of ELOF1, DSIF and NELF can counteract TFIIF, resulting in stable promoter-proximal paused RNA polymerase II.

Many factors in addition to DSIF-NELF-ELOF1 are associated with promoter-proximal pausing including TFIID, the +1 nucleosome, and FACT^11,60,77–80^. Our reconstituted experiments show that none of these factors are required for robust pausing in a minimal system. These factors may act as regulatory or accessory checkpoints that fine-tune the position or duration of promoter-proximal pausing induced by DSIF-NELF-ELOF1 in a chromatin context.

Our findings implicate ELOF1 in determining RNA polymerase II fate downstream of promoter-proximal pausing, including premature termination or release into productive elongation. Previous work has shown that ELOF1 loss slows release into productive elongation in cells^38,39,44,45^. The faster decay of paused RNA polymerase II we observe upon ELOF1 degradation is therefore unlikely to reflect accelerated pause release and instead may reflect increased premature termination. The Integrator complex can associate with the paused RNA polymerase II-DSIF-NELF complex and induce its termination. Structural studies of Integrator bound to RNA polymerase II-DSIF-NELF show dissociation of the DSIF DNA clamp, which includes SPT4 and SPT5 NGN and KOW1 domains, from RNA polymerase II to accommodate the Integrator tail module (INTS10-INTS13-INTS14-INTS15)^24^. Here, we show that part of the DSIF DNA clamp is stabilized by ELOF1, which may prevent engagement of Integrator tail module (Fig. 5F). The tail module is proposed to open the RNA polymerase II DNA clamp to promote dissociation of RNA polymerase II from DNA, and ELOF1 could potentially impede this activity by stabilizing the DNA clamp. Whether Integrator promotes premature termination in the presence of ELOF1 remains unknown. Finally, we show the DSIF-NELF-ELOF1 paused complex can be efficiently converted into EC* by P-TEFb and associated elongation factors, demonstrating that ELOF1-stabilized pausing is reversible. ELOF1 functionally appears to be like DSIF in the sense that it is critical for both pausing and productive elongation and would be retained after pause escape. Future work is needed to understand how ELOF1 toggles between pausing and elongation functions.

In summary, ELOF1 is a core component of the promoter-proximal paused complex and the long-sought TFIIF resistance factor. Together with its established role in transcription elongation, ELOF1 emerges as a central regulator of transcriptional output, contributing to both the establishment of pausing and the subsequent transition into productive elongation. control at pause sites could support the rate of release into transcription elongation or premature termination.

In summary, GATO-seq is a highly sensitive and modular technique that can allow for the study of all aspects of transcriptional regulation across species and will serve as a complement to cell-based approaches. We envision that GATO-seq will serve as a platform for bottom-up reconstitution of transcriptional processes. The GATO-seq approach, together with cell-based, biochemical, and biophysical assays, will enable a more complete and mechanistic understanding of gene regulation.

## Limitations of the study

ChIP-nexus samples were only collected for 3 time points after triptolide treatment. This may impact the robustness of the exponential decay model fit, and therefore, the number of genes affected by ELOF1degradation may be larger than reported. All biochemical experiments were performed on non-chromatinized DNA templates. The role of ELOF1 in transcriptional pausing may differ in a chromatin context. We observe no direct contact between ELOF1 and NELF in our cryo-EM reconstructions. This does not exclude an interaction with flexible regions of either protein.

## Declaration of Generative AI and AI-assisted technologies in the writing process

Claude 4.5-4.7 and ChatGPT 5.5 were used for copy-editing. All results were verified by the authors.

## Acknowledgments

We thank all past and present members of the Vos lab for support, discussion, and comments on the manuscript. We especially thank A. D’Ordine and T.J. Russell for their help in executing the triptolide experiments and V. Tholkes and A.J. Diao for initial cryo-EM sample preparation. Cryo-EM specimens were prepared and data was collected at the Cryo-EM Facility at MIT.nano, including use of the Talos Arctica gifted by the Arnold and Mabel Beckman Foundation. ChIP-nexus libraries were sequenced at the MIT BioMicro Center. We thank S. Sterling, J. Podgorski and D. Lim for support at MIT.nano, J. A. Marteijn for sharing HCT116 cell lines, D. Schatz, L. Wu, and A.D. Yadavalli for sharing RASH-1C cell lines and critical discussions, L. Farnung for sharing reagents and critical discussions, R. Lamason and H. Kurka Margolis for sharing reagents, and B. Pacheco-Fiallos for help with illustrations.

## Funding

National Institutes of Health grant DP2-GM146254 (SMV)

Freeman Hrabowski Scholar of the Howard Hughes Medical Institute (SMV)

Swiss National Science Foundation, Early Postdoc Mobility Fellowship (RVN)

Max Planck Gesellschaft (AM)

## Author contributions

Conceptualization: RVN,

SMV Methodology: RVN, SMV

Investigation: RVN, AB, ZS, AM, SMV

Visualization: RVN, AB, ZS

Funding acquisition: SMV, AM

Project administration: SMV

Supervision: SMV, AM

Writing – original draft: RVN, SMV

Writing – review & editing: RVN, AB, ZS, AM, SMV

## Lead contact

## Materials availability

Materials available upon reasonable request from Seychelle M. Vos.

## Data and code availability

Structure models coordinates are deposited in the PDB (XXXX) and the cryo-EM maps are deposited into the Electron Microscopy Data Bank (EMDB): Pol II-DSIF-NELF-ELOF1 composite map EMD-YYYY, Map A EMD-ZZZZ, Map B EMD-WWWW, and Map C EMD-VVVV

Oxford nanopore direct RNA sequencing and Illumina sequencing ChIP-nexus reads are available at SRA under the accession number PRJNAXXXXXXX and PRJNAYYYYYYY. Processed GATO-seq signal, and ChIP-nexus data is available at the GEO under the accession number GSEZZZZZZ.

## Methods

### Experimental model and subject details

Bacterial *E. coli* strains DH5?, BL21 (DE3) RIL (Merck), DH10aEMBacY (Geneva Biotech) were grown in standard LB or 2xYT media at 37 ºC supplemented with ampicillin (100 µg/ml), gentamycin (10 µg/ml), kanamycin (30 µg/ml), or chloramphenicol (34 µg/ml).

Insect cells Sf9 (Expression Systems), Sf21 (Expression Systems), and Hi5 (Expression Systems) cells were cultured in ESF 921 media (Expression Systems) at 27ºC.

Porcine (*Sus scrofa*) thymus tissue was obtained from Pel-Freez Biologicals. The tissues were obtained from young and healthy pigs, frozen at -20ºC, and stored at -80ºC prior to use.

HCT116 Human colorectal cancer cell line was grown adherent in DMEM media (Gibco) supplemented with 10 % FBS (Gibco) and 0.5 mg/mL penicillin-streptomycin-glutamine at 37 °C and 5 % CO_2_. RASH-1 cells were grown in suspension in RPMI media (Gibco), supplemented with 10 % FBS (Gibco) and 0.5 mg/mL penicillin-streptomycin at 37 °C and 5 % CO_2_.

## METHODS DETAILS

### Cloning and protein expression

Transcription initiation factors TFIIA, B, E, F, H, CAK, and TBP were cloned and expressed as previously described^54^. Elongation factors DSIF^81^, NELF^82^, ELOF1^43^, TFIIS^59^, P-TEFb, PAF1c, RTF1, and SPT6^16,55^ were cloned and expressed as detailed previously.

### Protein Purification

*S. scrofa* RNA polymerase II was purified as previously described from thymus tissue^58,59,83,84^.

DSIF was purified as previously described^81^, employing a nickel affinity step, (HisTrap HP column (Cytiva)), 3C protease affinity tag cleavage, an anion exchange purification step (HiTrap Q column (Cytiva)) and size exclusion chromatography (HiLoad S200 16/600 pg column (Cytiva)).

NELF was purified as previously described^82^, using a nickel affinity step in series with an anion exchange step, followed by TEV protease affinity tag cleavage. After removing uncleaved protein, the sample was further purified by size exclusion chromatography (HiLoad S200 16/600 pg column (Cytiva)).

ELOF1 was purified as previously described^43^, using nickel affinity purification, followed by TEV protease affinity tag cleavage, removal of uncleaved products by nickel affinity purification and finally purified by size exclusion chromatography (HiLoad S75 16/600 pg column (Cytiva)).

Initiation factors TFIIA, B, E, F, H, CAK, TBP^54,85^, and elongation factors TFIIS, PAF1c, RTF1, P-TEFb and SPT6^16,55^ were purified as described previously.

### RNA extension assays

DNA and RNA oligos were synthesized and purchased from Sigma-Aldrich, resuspended in water to a final concentration of 100 µM, aliquoted, flash-frozen in liquid nitrogen, and stored at −80 °C. Transcription assays were performed with perfectly complementary scaffolds; the 9-base-pair DNA– RNA hybrid is preceded by 13 nucleotides of upstream DNA, and preceded by 28 nucleotides of downstream DNA. 6 nucleotides of exiting RNA are coupled to a 5′-6-FAM label. Assays reported with bacterial elemental pause sequence were designed as previously described using the following sequence: template DNA 5′-CCA CTG GAA GAT CTG AAT TTA CGG GCG CAA CTA TGC CGG ACG TAC TGA CC -3′, non-template DNA 5′-GGT CAG TAC GTC CGG CAT AGT TGC GCC CGT AAA TTC AGA TCT TCC AGT GG -3′, RNA 5′-6-FAM-UUU UUU GGC AUA GUU-3′. RNA and template DNA were mixed in equimolar ratios and were annealed by incubating the nucleic acids at 95 °C for 5 min and then decreasing the temperature by 1°C per minute steps to a final temperature of 30 °C in a thermocycler in a buffer containing 100 mM NaCl, 20 mM Na-HEPES pH 7.4, 3 mM MgCl_2_, and 10% (v/v) glycerol. All concentrations correspond to the final concentrations used in the assay. *S. scrofa* RNA polymerase II (100 nM) and the RNA-DNA template hybrid (100 nM) were incubated for 10 min at 30 °C, shaking at 300 rpm. Next, the non-template DNA strand was added (100 nM) and incubated for another 10 min. Factors were diluted in protein dilution buffer (300 mM NaCl, 20 mM Na-HEPES pH 7.4, 10% (v/v) glycerol and 1 mM DTT) and mixed to a final concentration of 300 nM. The reactions were then diluted to achieve final assay conditions of 100 mM NaCl, 20 mM Na-HEPES pH 7.4, 3 mM MgCl_2_, 4% (v/v) glycerol, 1 mM DTT, 25 µM ZnCl_2_ and further incubated for 10 min at 30 °C. Transcription reactions were initiated by adding NTPs at a final concentration of 10 µM. Reactions (5 µL) were quenched after 0-10 min in 5 µL 2x Stop buffer (6.4 M urea, 50 mM EDTA pH 8.0, 1x TBE buffer and 4 µg of proteinase K (New England Biolabs)). Quenched samples were incubated for 30 min at 37°C and RNA products were separated by denaturing gel electrophoresis (5 µL of sample applied to an 8 M urea, 1x TBE, 20% Bis-Tris acrylamide 19:1 gel run in 0.2x TBE buffer at 300 V for 160 min). Products were visualized using a FAM filter on a Typhoon 9500 FLA Imager (GE Healthcare Life Sciences) at 900 PMT. Gel images were quantified using ImageJ (v1.53K). A 0.48 × 1.98 cm box surrounding each lane was created to obtain lane plots. A baseline was manually placed, and the tracing tool was used to obtain the total integrated density for each band. The integrated density of the +7 band (full-length product) was normalized relative to the intensity of the starting RNA band, by dividing the full-length band integrated density value by that of the starting RNA band. Graphs were prepared in GraphPad Prism 10. Each point represents the mean intensity from three individual replicates.

### Library of templates for GATO-seq

GATO-seq template library was generated as previously described^54^. Briefly, the coordinates of the top 1,000 human promoter-proximal sequences (hg38) ranked by transcription start site (TSS) focus score derived from PRO-seq and RNA-seq data were used. Genomic regions spanning the TSS to +261 bp were retrieved and synthesized as an oligonucleotide pool (Twist Biosciences) with flanking cloning adaptors. The pool was amplified and cloned upstream of an AdML promoter in a pBlueScript plasmid vector by Gibson assembly, transformed into *E. coli*, and amplified to generate the plasmid library. Linear DNA templates for transcription assays were produced by emulsion PCR and purified by anion-exchange chromatography. Synthetic RNA spike-ins of defined lengths were generated or synthesized to calibrate sequencing length biases.

### GATO-seq sample preparation and analysis

GATO-seq experiments were performed as described^54^. Preinitiation complexes were assembled stepwise with purified RNA polymerase II and general transcription factors on the promoter library templates, followed by controlled transcription initiation and elongation reactions under defined nucleotide concentrations (10 µM or 1 mM). Reactions were quenched; RNA products were purified, polyadenylated, and prepared for Oxford Nanopore direct RNA sequencing. Sequencing libraries were generated using the SQK-RNA004 kit and run on MinION flow cells. Reads were basecalled using Dorado 10 (Oxford Nanopore Technologies) super high accuracy model, aligned to the reference library and spike-in sequences with Bowtie2 (v2.4.5)^86^, and processed to extract RNA 3′ ends with deepTools (c3.5.5)^87^. Spike-in normalization was used to correct length-dependent sequencing bias, and normalized signals were used to compute transcription passthrough indices, identify pause sites, and analyze pause dynamics using custom Python scripts and deepTools. Full experimental and computational details are described in ^54^.

### Cryo-EM sample preparation

The consensus super pause sequence described before was used containing the following sequences: DNA 5′-CCA CTG GAA GAT CTG AAT TTG CGG CAG CAG CTC CGC CGG ACG TAC TGA CC -3′, non-template DNA 5′-GGT CAG TAC GTC CGG CGG AGC TGC TGC CGC AAA TTC AGA TCT TCC AGT GG -3′, RNA 5′-6-FAM-UUU UUU GGC GGA GCU-3′.

The RNA polymerase II complex was assembled as described above for the RNA extension assay but at a higher concentration (1 µM instead of 100 nM final). RNA polymerase II (100 pmol) was mixed with 125 pmol RNA-DNA template, followed by 125 pmol non-template DNA. The final buffer condition consisted of 100 mM NaCl, 20 mM Na-HEPES pH 7.4, 3 mM MgCl_2_, 1 mM DTT, and 4% (v/v) glycerol. The sample was incubated for 30 min at 30°C. RNA extension was enabled by adding CTP, GTP and UTP to a final concentration of 10 µM and incubated for 5 min at 30 °C. The reaction was quenched with EDTA pH 8.0 at a final concentration of 10 mM and transferred to a 4 °C ice bath. RNA polymerase II-DSIF-NELF-ELOF1 complex was purified by applying it to a Superose 6 increase 3.2/300 column equilibrated in buffer matching reaction conditions (100 mM NaCl, 20 mM Na-HEPES pH 7.4, 4% (v/v) glycerol, 3 mM MgCl2, and 1 mM DTT) on an Äkta Micro purification system (Cytiva) at 4°C. 50 µL fractions were collected and analyzed by SDS-PAGE followed by Coomassie staining and TBE-urea gel electrophoresis.

The peak fractions containing RNA polymerase II, DSIF, NELF, ELOF1 and nucleic acids were crosslinked with a final concentration of 0.1% (v/v) glutaraldehyde for 10 min on ice and quenched with 8 mM aspartate and 2 mM lysine. The crosslinked sample was dialyzed for 4 hours at 4°C against a buffer containing 100 mM NaCl, 20 mM Na-HEPES pH 7.4, 1 mM DTT, and 3 mM MgCl_2_ in 10 kDa MWCO Slide-A-Lyzer MINI Dialysis Units (Thermo-Fisher). Dialyzed samples at a final concentration of 300-400 nM were applied to R2/2 UltrAuFoil grids (Quantifoil). The grids were glow-discharged for 2 min before applying 4 µL of sample to each side of the grid (8 µL total), incubated for 4 s, and blotted for 4s. Vitrification was done by plunging the grids into liquid ethane using a Vitrobot Mark IV (FEI Company) operated at 4°C and 100% humidity.

### Cryo-EM data collection and processing

Micrographs were collected on a FEI Titan Krios II transmission electron microscope operated at 300 keV. A K3 summit direct detector (Gatan) with a BioQuantum energy filter (Gatan) was operated with a slit width of 20 eV. Data were automatically acquired at a nominal magnification of 105,000× using EPU (FEI). The corresponding pixel size was 0.822 Å per pixel. 50 frame image stacks were collected over 1.66s in counting mode at a dose flux of 20.4 e^−^/pix/s resulting in a total dose of 50.1 e^−^/Å^2^. A total of 33,437 image stacks were collected.

Movies were processed in CryoSPARC live v5.0.2 ^88^. The frames were motion-corrected and the contrast-transfer-function was estimated. Particles were then auto-picked using the blob picker tool and extracted using a box size of 400 pixels resulting in a total of 8,251,315 particles. Particles were subjected to 2D classification and classes containing secondary structure features were selected for ab-initio 3D reconstruction. A second ab-initio job was performed using 2D classes with non-aligning particles. All particles were subjected to heterogeneous refinement using the ab-initio volumes as input references to generate 6 3D classes. 1,764,508 particles showed features for RNA polymerase II bound to DSIF, NELF and ELOF1, and these particles were refined to 2.6 Å (FSC 0.143). Particles were further classified to sort for compositional and conformational heterogeneity as described below using CryoSPARC v5.0.2.

Initial focused classifications were performed to separate particles containing NELF from those lacking NELF. Particles containing NELF in the paused conformation were further classified to identify particles with densities corresponding to ELOF1 and SPT5 NGN-SPT4. This set of particles was refined and used as an input volume in a heterogeneous refinement resulting in 398,080 particles with strong densities corresponding to NELF in the paused conformation.

After homogeneous refinement, a further round of 3D classification was performed on this set of particles to identify particles with strong occupancy for ELOF1, SPT4, SPT5 NGN, SPT5 KOW1 and the upstream DNA resulting in 187,498 particles and a map with an overall resolution of 2.8 Å (FSC 0.143) (Map A). Finally, separate focused classifications were performed to enhance densities for NELF-BE and the upstream DNA-SPT5 KOW1. The particle sets from these two classifications were individually subjected to reference-based motion correction, Global CTF correction, and homogeneous refinement. The NELF-BE focused map contained 88,105 particles and refined to 2.7 Å (FSC 0.143) (Map B). The upstream DNA-SPT5 KOW1 focused map contained 83,871 particles and refined to 2.8 Å (FSC 0.143) (Map C). A composite map was generated from the refined half-maps using FrankenMap in the Warp suite^89^.

Our initial classification unveiled particles that lacked NELF or had NELF in the poised conformation. Particles lacking NELF were subjected to focused 3D classifications with a soft mask to identify particles with densities corresponding to DSIF, ELOF1, or both DSIF and ELOF1. Maps lacking density for DSIF exhibited mobile RPB1 clamp densities. Focused classifications on the RPB1 clamp domain were performed to identify particles with stable clamp domains that were engaged with DNA. Maps containing densities for DSIF or DSIF-ELOF1 were further classified on their active sites to define the translocation state of RNA polymerase II. A subset of 62,930 particles showed a sidetracked register and were subjected to homogeneous refinement to a 2.7 Å (FSC 0.143) map (Map D). Poised NELF particles were further classified to improve the densities for NELF-BE and subjected to homogeneous refinement.

For focused classifications, masks were generated using published models (PDB 8UHD, 8UHG and 9BZ0). Masks were generated from low-pass filtered volumes (15 Å) that were dilated by 12 Å and extended by 5 soft pixels. Active site classifications were performed with 10 Å low filtered masks that contained the bridge helix, trigger loop, and the first 3 bases of the RNA DNA hybrid. The mask was extended by 3 pixels and 1 soft pixel. All final maps were sharpened using a B-factor of -15 Å^2^.

### Cryo-EM model building

Previously published models were initially rigid body docked into our cryo-EM maps using UCSF ChimeraX (v1.7). RNA polymerase II and ELOF1 from PDB 9BZ0 were rigid body fit in the refined DSIF-NELF-ELOF1 NELF-BE focused cryo-EM map (Map B). Nucleic acids, NELF, SPT5 KOW1-5 (residues 276 to 753) from PDB 8UHD and SPT4 and SPT5’s NGN domain (residues 176 to 266) from PDB 3H7H were rigid body fit into the cryo-EM composite map. The upstream DNA duplex encompassing 8 bp was rigid body fit into a 6Å low-pass filtered volume from the DSIF-NELFELOF1 upstream DNA focused map (Map C). Four additional deoxynucleotides of the non-template DNA strand adjacent to the upstream DNA duplex were modelled using this low-pass filtered map. Interestingly, additional density near the RPB2 lobe residue Y262 and -9 K334, K337, K340 was observed. This density was present in all maps, except RNA polymerase II lacking factors. This density is not clearly connected to densities associated with other elongation factors, and thus the source of this signal remains to be determined. Densities for RPB4 and RPB7, SPT5’s KOW1 to x-4 were weak and were not adjusted after the rigid body fit. The first 15 residues of ELOF1 were not visible, as well as RPB1 Trigger loop residues 1103-1115 and therefore were not included in the final model. Because we used actively transcribing complexes, unambiguous assignment of the nucleic acid sequence register was not possible. The DNA and RNA were modeled by substituting the sequence from PDB 8UHD to match the assumed state with 3-nucleotide RNA extension by RNA polymerase II. 15 bp of the down-stream DNA duplex, 9 bp of the RNA-DNA hybrid and 12 bp of upstream DNA duplex were modelled. Density for eight deoxynucleotides of the non-template DNA strand were not visible and therefore were not included in the final model. Side chains were manually adjusted in Coot 1.1.08 113^90^ using the cryo-EM composite map and maps B and C. Input models for refinement were prepared using ReadySet in Phenix. Final models were real space refined in Phenix version 1.21.2 114^91^ with secondary structure and Ramachandran restraints. Model statistics and data collection information are provided in Table 1.

### ChIP-nexus

ChIP nexus was performed following the procedure originally described^47^with the modifications described here: https://stowers-institute.files.svdcdn.com/production/files/20210812_ChIP-nexus-protocol-1.pdf?dm=1732108008. HCT116 cells were seeded in 150 mm culture dishes and grown to 90 % confluency (50 million cells). Adherent cells were washed two times with 15 mL ice-cold PBS buffer and crosslinked with 1% (v/v) Pierce methanol-free formaldehyde (Thermo Fisher) at room temperature for 10 minutes with gentle shaking. The crosslinking reaction was quenched with 2 mM glycine at room temperature for 5 minutes. Cells were washed three times with ice-cold PBS supplemented with EDTA-free protease inhibitors (Roche). Cells were scraped from the dish and resuspended in 15 mL ice-cold PBS with protease inhibitors. Resuspended cells were counted and aliquoted in 10 million cell batches. Cells were spun, buffer was removed, and cell pellets were flash frozen in liquid nitrogen and stored at -80 °C until use.

RASH-1 cells were seeded and grown to a final density of 1 million cells/mL. Cells were treated with dTAGV-1 (Tocris, 6914) or dTAG-NEG (Tocris, 6915) at a final concentration of 0.5 µM for 3 hours at 37 °C. After treatment, cells were washed twice with ice-cold PBS, counted and aliquoted in 50 million cell batches. Each batch was crosslinked with 1 % (v/v) Pierce methanol-free formaldehyde (Thermo Fisher) at room temperature for 10 minutes with gentle shaking and quenched with 2 mM glycine at room temperature for 5 minutes. Cells were washed twice with 15 mL ice-cold PBS buffer supplemented with 1X protease inhibitors (Roche). Cell pellets were directly used for immunoprecipitation.

For triptolide treatment experiments, RASH-1 cells treated with dTAGV-1 or dTAG-NEG for 3 hours were further treated with 10 µM Triptolide (Sigma) or DMSO vehicle and harvested by centrifugation at the indicated experimental times. The samples were then processed as described above. Cell pellets were resuspended in 130 µL ChIP buffer (15 mM HEPES pH 7.5, 140 mM NaCl, 1 mM EDTA, 0.5 mM EGTA, 1% (v/v) Triton X-100, 0.5% (w/v) N-lauroylsarcosine, 0.1% (w/v) sodium deoxycholate, 0.1% (w/v) SDS, with freshly added protease inhibitors). Sonication was performed with a CovarisE220 focused-ultrasonicator for 10 minutes at a peak power = 105, duty factor = 2 and 200 cycles. Chromatin extracts were then centrifuged at 16000 x g for 30 min at 4°C, and supernatants were used for ChIP.

To couple Dynabeads with antibodies, 50 µL Protein G Dynabeads (Thermo Fisher) were used for each ChIP-nexus experiment and washed twice with ChIP Buffer. Dynabeads were resuspended in 400 µL ChIP Buffer, and 8 µg antibodies were added: anti-RPB1 (D8L4Y, Cell Signaling Technology), anti-SPT5 (611107, BD Biosciences), and anti-HA (C29F4, Cell Signaling Technology) along with 4 µg spike-in antibody (61686 Active Motif). Tubes were incubated at 4°C for 2 hours with rotation. After the incubation, antibody-bound beads were washed twice with ChIP Buffer.

Chromatin extracts were combined with antibody-coupled beads and 240 ng spike-in chromatin (53083 Active Motif) was added to each condition and incubated at 4°C overnight with rotation. Beads were washed between each enzymatic treatment with Nexus washing buffer A to D (wash buffer A: 10 mM Tris-EDTA pH 8.0, 0.1% (v/v) Triton X-100, wash buffer B: 150 mM NaCl, 20 mM Tris-HCl, pH 8.0, 5 mM EDTA, 5.2% (w/v) sucrose, 1.0% (v/v) Triton X-100, 0.2% (w/v) SDS, wash buffer C: 250 mM NaCl, 5 mM Tris-HCl pH 8.0, 25 mM HEPES pH 7.4, 0.5% (v/v) Triton X-100, 0.05% (w/v) sodium deoxycholate, 0.5 mM EDTA pH 8.0, wash buffer D: 250 mM LiCl, 0.5% (v/v) IGEPAL CA-630, 10 mM Tris-HCl pH 8.0, 0.5% (w/v) sodium deoxycholate, 10 mM EDTA pH 8.0) and rinsed with 10 mM Tris-HCl pH 8.0. End repair and dA-tailing were performed using the NEBNext End Repair Module and the NEBNext dA-Tailing Module (NEB). ChIP-nexus adaptors with mixed fixed barcodes (CTGA, TGAC, GACT, ACTG) were ligated with Quick T4 DNA ligase (NEB) and converted to blunt ends with Klenow fragment and T4 DNA polymerase (NEB). The samples were treated with lambda exonuclease (NEB). Chromatin was eluted from the antibodies by an overnight incubation at 65 °C in 25 mM Tris pH 8.0, 5 mM EDTA pH 8.0 and 0.5% (w/v) SDS. DNA was purified by phenol-chloroform extraction and ethanol precipitation. Purified single-stranded DNA was then circularized with CircLigase (Lucigen) and subjected to PCR amplification with NEBNext High-Fidelity Q5 2X PCR Master Mix and ChIP-nexus primers containing sequencing indexes. The ChIP-nexus libraries were then gel-purified before high-throughput sequencing by Illumina NovaSeq.

### ChIP-nexus data processing

UMIs (barcode pattern NNNNNXXXX) were extracted from ChIP-nexus reads with UMI-tools^92^ and written to read tags. Reads were trimmed with Trimmomatic v0.39^93^ using the parameters -phred33, SLIDINGWINDOW:4:15, ILLUMINACLIP:<adaptors_fasta>:2:30:10, LEADING:3, TRAILING:3, and MINLEN:36; Illumina single-ended adapter sequences were supplied. Trimmed reads were aligned to the human reference genome (GRCh38.p12) with Bowtie2 v2.3.5.1^86^ using --no-mixed, --no-unal, and --phred33. Alignments were deduplicated in a UMI-aware manner with UMICollapse^94^. Protein–DNA binding sites were called with MACS2 v2.2.7.1^95^ using callpeak with the options --nolambda, --nomodel, --keep-dup all, --extsize 200, -B, --SPMR, -f BAM, and -g hs.

To generate metagene profiles, we separated forward- and reverse-strand alignments with SAMtools view^96^ (-F 16 for forward, -f 16 for reverse) and derived 5′-end coverage with BEDTools^97^ genomecov (-5 -bg). Coverage tracks were then normalized by applying the sample-specific background scaling factors estimated by DiffBind v3.0.15^98^ (--scaleFactor).

### ChIP-nexus differential binding analysis

To identify protein–DNA binding sites with altered occupancy across conditions, we used DiffBind, an R package for filtering artifacts, generating consensus peaks, quantifying signal, normalizing counts, and testing for differential binding. Signals overlapping ENCODE blacklist regions of the human hg38 genome were removed with dba.blacklist (blacklist=DBA_BLACKLIST_HG38). Peaks across samples were merged into consensus peaks and quantified using dba.count with minOverlap=2, fragmentSize=0, summits=0, bRemoveDuplicates=FALSE, and bSubControl=FALSE. Size factors for statistical testing were computed with dba.normalize (normalize=DBA_NORM_RLE, background=TRUE). Differential binding sites were then identified with the standard DESeq2^99^ pipeline using the precomputed counts and size factors from DiffBind.

#### Pol II Half-life Estimation

To estimate Pol II half-life in promoter-proximal regions (TSS-0.1Kb to TSS+0.5Kb), we fit an exponential decay model, N(t) = N0 · e−αt, to Pol II occupancy measured at 0, 5, and 20 minutes after triptolide treatment using nonlinear least squares (nls) in R. Initial values were set to N0 = y0 (mean occupancy at 0 minute) and α0 = −2 · log(2) / y0. We constrained the fitted decay rate to α ≥ 0.023. Half-life was calculated as t1/2 = ln(2) / α. Regions were excluded if they were removed by independent filtering (using base-Mean as the filter statistic) in the corresponding DESeq2 analysis, or if they showed <20% reduction after 20 minutes of triptolide treatment.

