## Supplementary figures for "ELOF1 is a core component of the promoter-proximal paused RNA polymerase II complex"

#### Figure S1

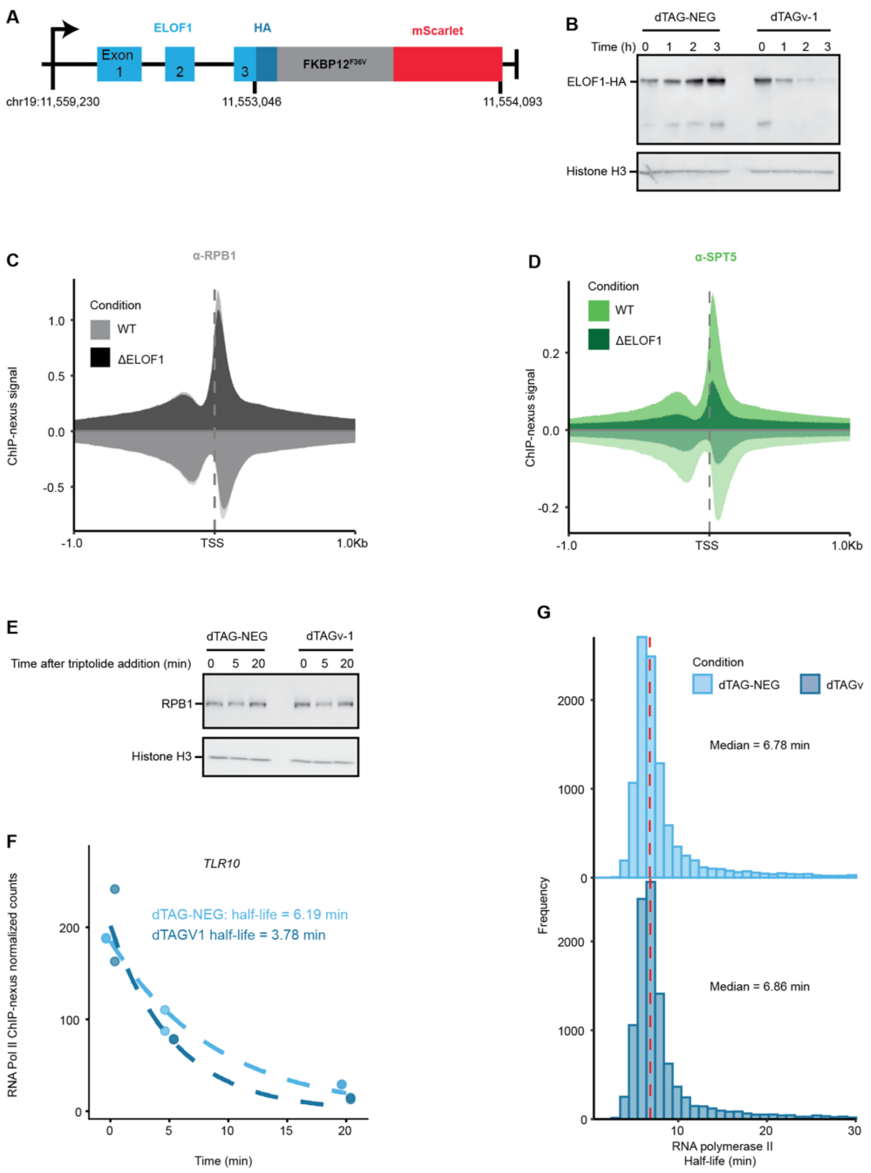


**Supplementary Figure 1. ChIP-nexus in ΔELOF1 cells, quality control of ELOF1 degradation, and RNA polymerase II half-life estimation.**

**A)** Schematic of ELOF1 endogenous gene fused to HA tag, degron tag and mScarlet as reported in[44] **B)** Western blot using an anti-HA antibody in ELOF1-degron RASH-1C cells after treatment with dTAG-NEG or dTAGV-1 at the indicated time points. **C)** Normalized ChIP-nexus metagene profiles from HCT116 cells parental (WT) and ELOF1 homozygous knockout

(ΔELOF1), using an antibody directed against the RPB1 N-terminus. The positive strand signal is shown above the baseline (darker shade) and the negative strand signal below the baseline (light shade). Average trace from two biological replicates. **D)** Normalized ChIP-nexus metagene profiles as in A) using an anti-SPT5 antibody. Average trace from two biological replicates. **E)** Western blot using an anti-RPB1 CTD antibody (8WG16) in ELOF1-degron RASH-1C cells after 3 hours of treatment with dTAG-NEG or dTAGV-1, followed by the indicated time points after treatment with triptolide. **F)** Representative gene example (*TLR10*) with half-lives of promoter-proximal RNA polymerase II (TSS-0.1Kb to TSS+0.5Kb) after treatment with dTAG-NEG or dTAGV-1. The signal for RNA polymerase II decays upon triptolide treatment and was fit to an exponential decay model (dotted line). Values from two biological replicates are shown. **G**) Distribution of RNA polymerase II promoter-proximal half-life after treatment with dTAG-NEG (top) or dTAGV-1 (bottom). Red line indicates median value.

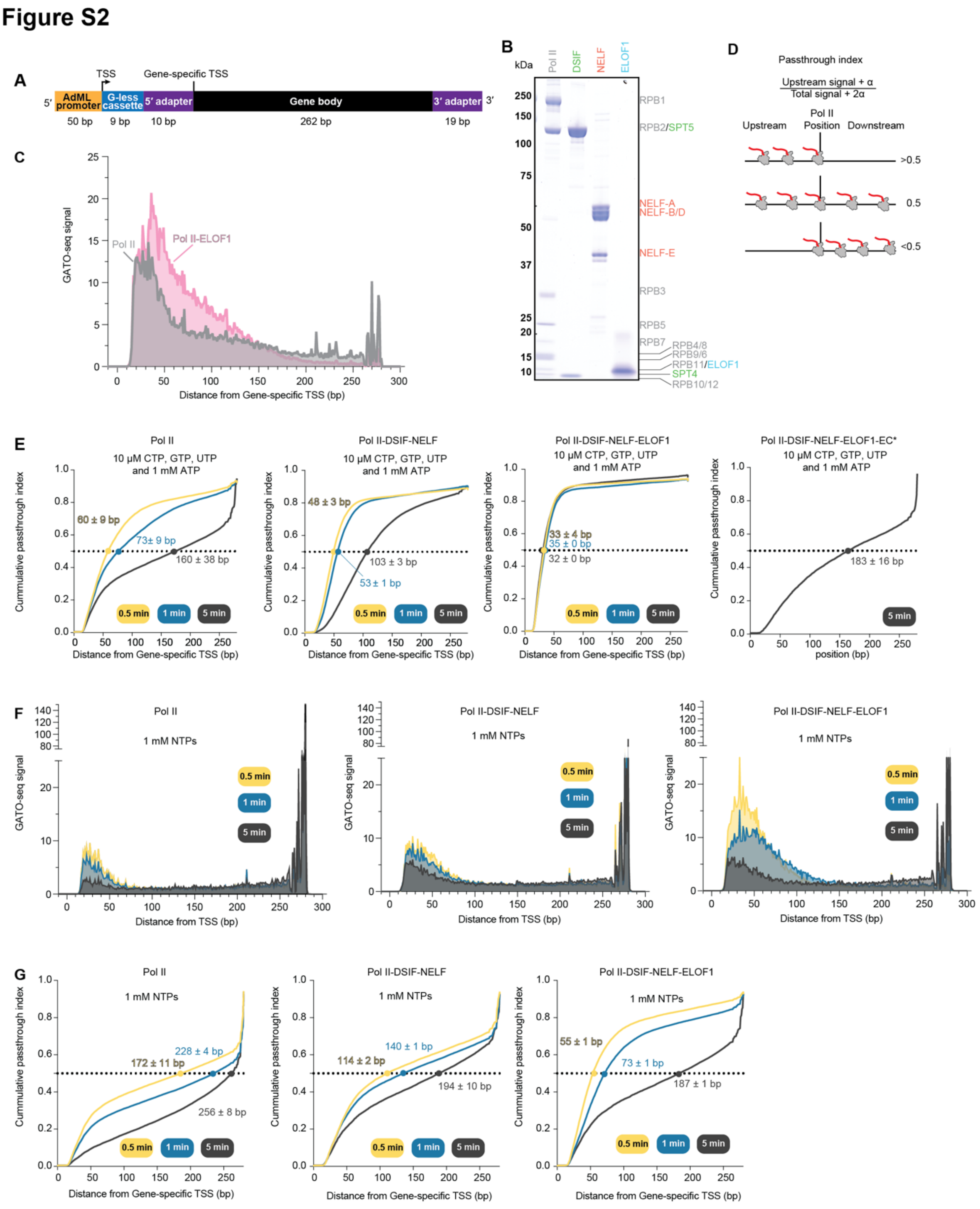


**Supplementary Figure 2. GATO-seq library composition, protein preparation, and effect of ELOF1 on GATO-seq reactions.**

**A)** Schematic of DNA template design employed in GATO-seq library. **B)** Coomassie stained SDS-PAGE (4-12%) gels of purified proteins (RNA polymerase II 2 µg, DSIF 0.8 µg, NELF 1.5 µg and ELOF1 0.25 µg.) **C)** Metagene profiles of GATO-seq signal in the absence and presence of ELOF1 after 1 minute from two biological replicates employing 10 µM CTP, GTP, UTP and 1 mM ATP. **D)** Schematic of passthrough index estimation. **E)** Comparison of passthrough index cumulative distributions at three different time points from two biological replicates with 10 µM CTP, GTP, UTP and 1 mM ATP and in the absence of elongation factors. (Pol II)[54], with DSIF and NELF[54], with DSIF, NELF and ELOF1, and at 5 minutes with DSIF, NELF, ELOF1 and EC* factors. Circles indicate cumulative distribution midpoint (y=0.5). **(F)** Metagene profiles of GATO-seq signal at three different time points from two biological replicates employing 1 mM NTPs, and in the absence of elongation factors (Pol II)[54], with DSIF and NELF and with DSIF, NELF and ELOF1. Average signal from two biological replicates. **(G)** Comparison of passthrough index cumulative distributions as in E) but with 1 mM NTPs and in the absence of elongation factors. (Pol II)[54], with DSIF and NELF, and with DSIF, NELF and ELOF1.

### Figure S3

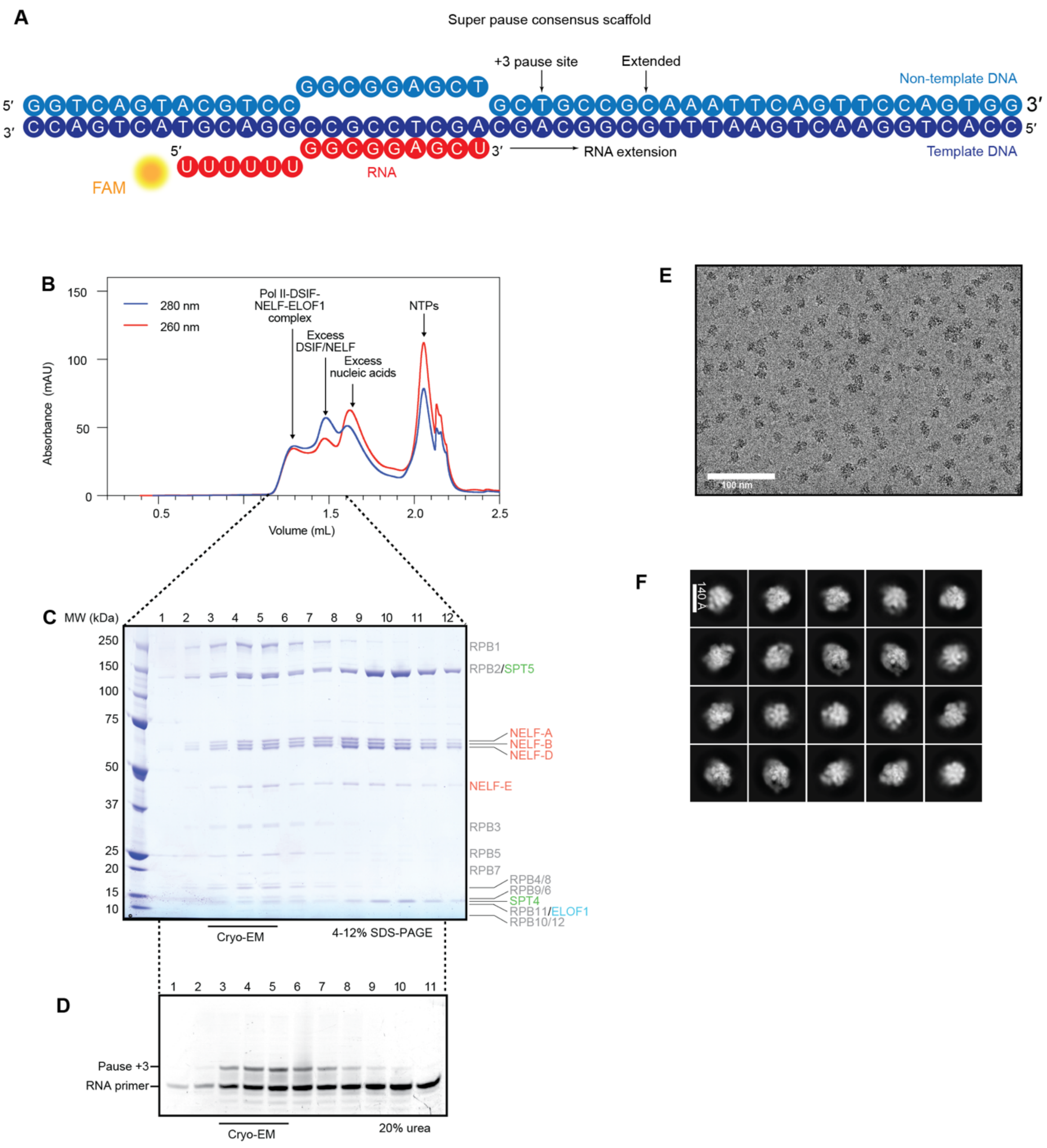


**Supplementary Figure 3. Sample composition and preparation of samples for cryo-EM and initial cryo-EM processing.**

**A)** Super pause consensus sequence scaffold used for RNA extension assays. **B)** Size exclusion chromatography trace of transcribing RNA polymerase II-DSIF-NELF-ELOF1 complex on

Super pause scaffold. **C)** Coomassie stained SDS-PAGE of fractions from chromatogram in A). **D)** 6 M urea, 20% polyacrylamide gel of fractions from (A) scanned in the FITC channel. **E)** Representative cryo-EM micrograph at -2.5 µm defocus. Scale bar shown on bottom left. **F)** Representative 2D classification images of RNA polymerase II-DSIF-NELF-ELOF1 particles. Scale bar is shown on the top left.

### Figure S4

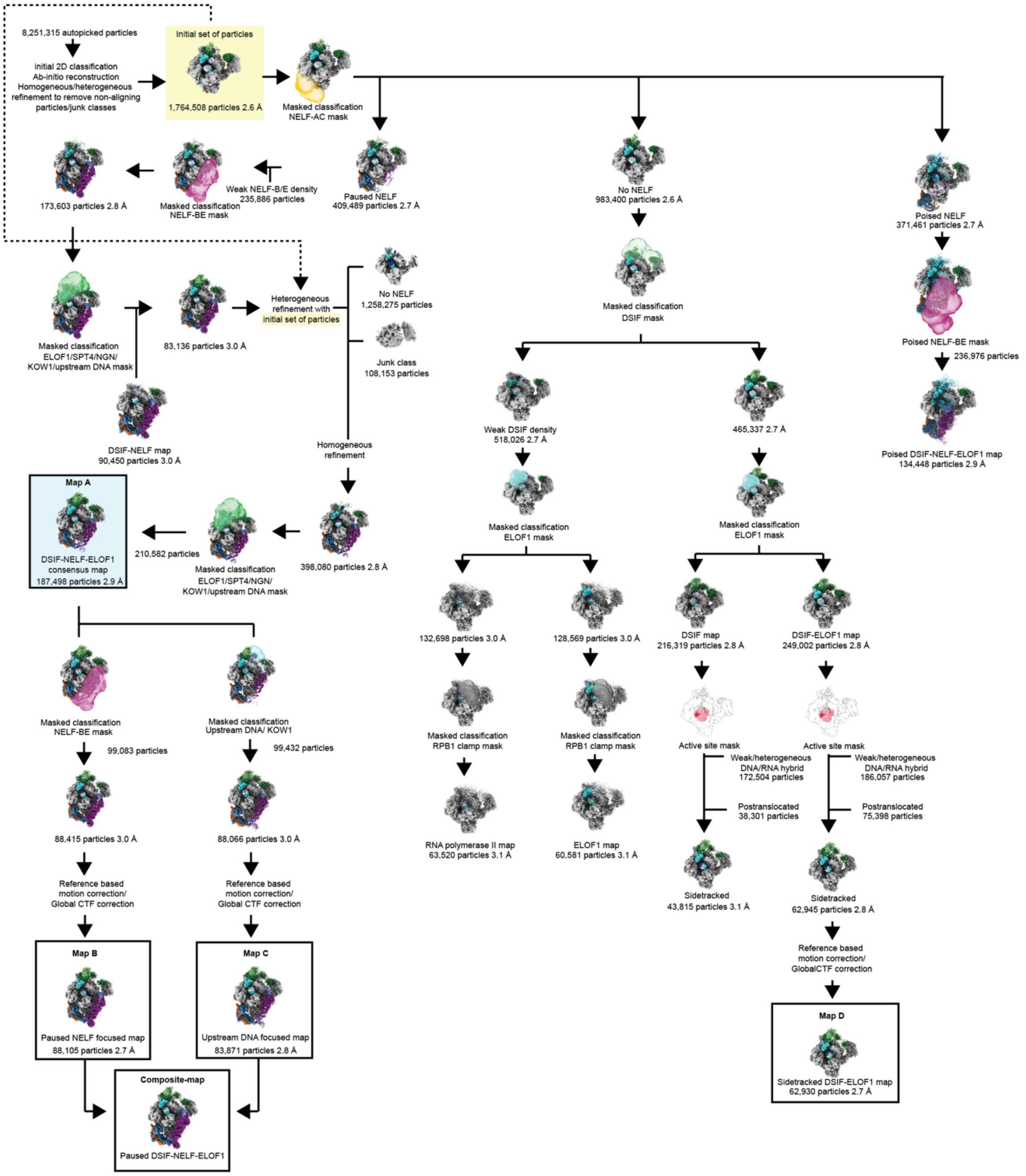


**Supplementary Figure 4. Cryo-EM processing tree.**

Cryo-EM classification tree for RNA polymerase II-DSIF-NELF-ELOF1 complex. Resolutions are shown for maps used for further classification. Masks used for focused classification are shown.

### Figure S5

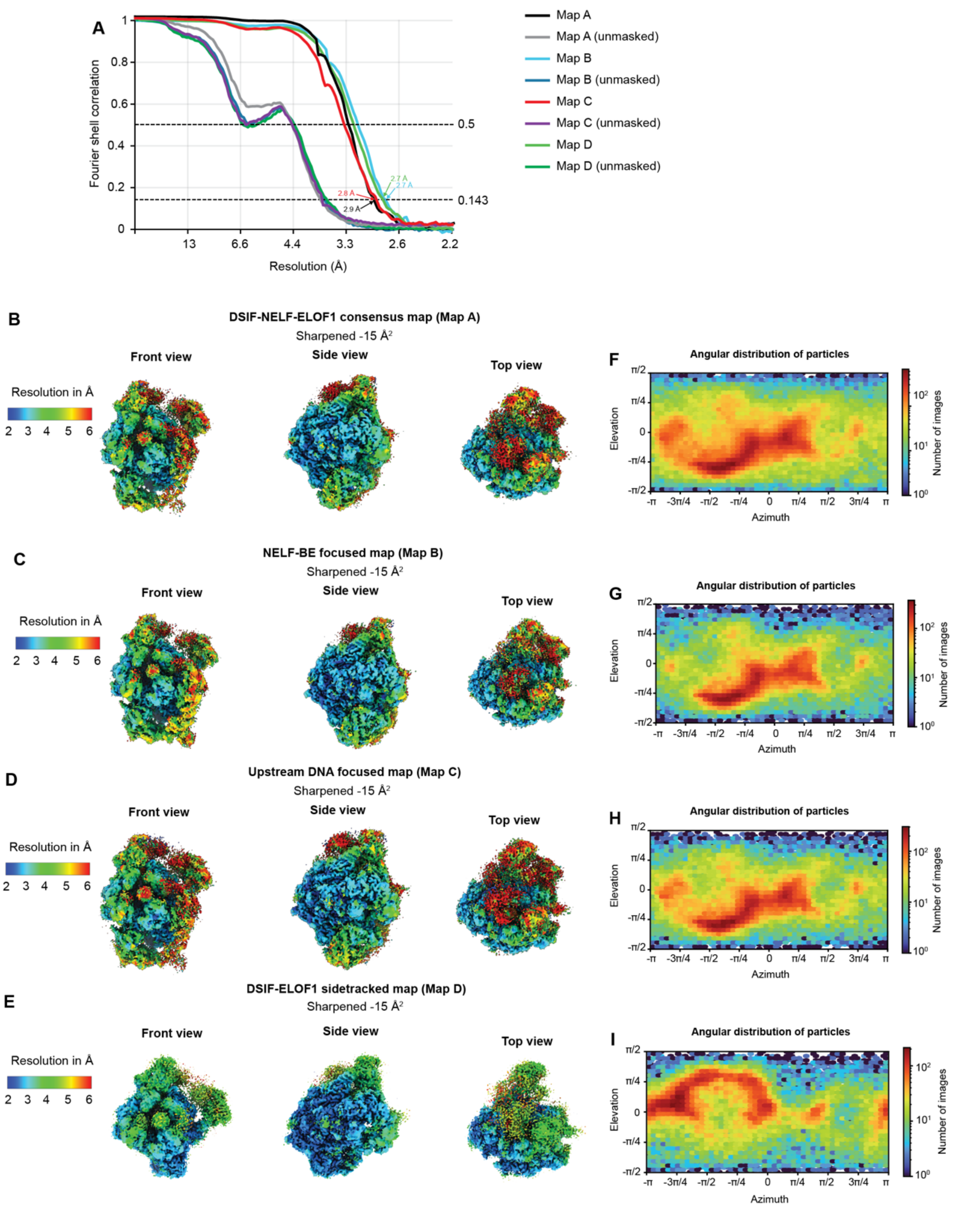


**Supplementary Figure 5. Cryo-EM data quality. A)** Fourier Shell Correlation curves (corrected and unmasked) of DSIF-NELF-ELOF1 consensus, upstream DNA focused, NELF-BE focused and DSIF-ELOF1 maps. FSC 0.5 and gold standard 0.143 are shown as horizontal dashed lines. Map resolutions are indicated. **B), C), D), E)** Cryo-EM reconstructions colored by local resolution (left), shading scale shown on left of B) Map A, C) Map B, D) Map C, E) Map D. **F), G), H), I)** Azimuth graphs of angular distribution of particles (right) from overall refinements for F) Map A, G) Map B, H) Map C, I) Map D.

### Figure S6

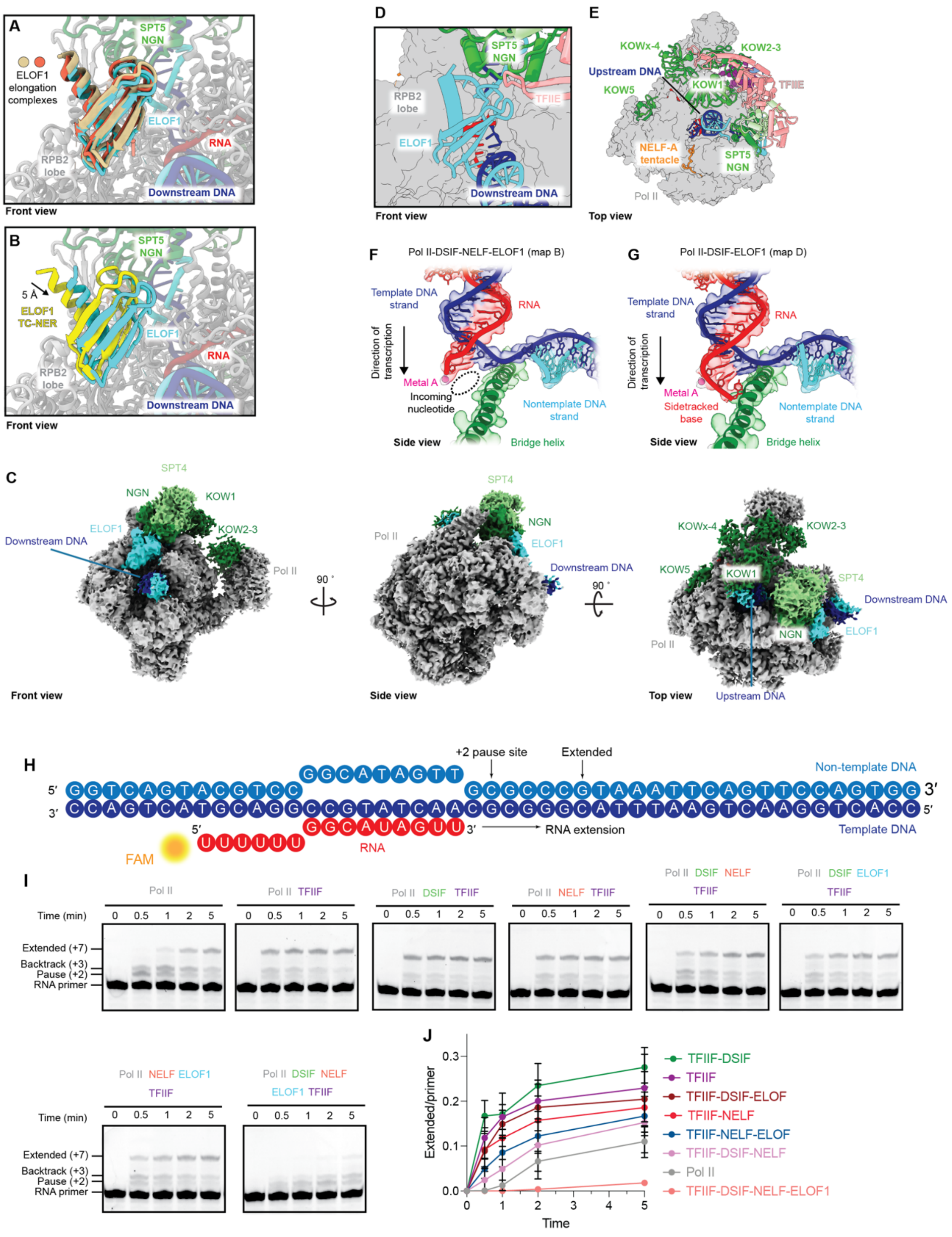


**Supplementary Figure 6. Comparison of RNA polymerase II-DSIF-NELF-ELOF1 with other complexes and DSIF-NELF-ELOF1 competition with TFIIF.**

**A)** Overlay of ELOF1 structures from activated elongation complexes (PDB 8XRM wheat color and PDB 9MLC orange color). **B)** Overlay of ELOF1 structures as in A) but with ELOF1 from TC-NER RNA polymerase II complex (PDB 9BZ0 in yellow color) **C)** Paused RNA polymerase II-DSIF-ELOF1 complex cryo-EM map. RNA polymerase II is colored silver, template DNA blue, non-template DNA cyan, and RNA red. SPT4 is colored light green and SPT5 dark green. ELOF1 is shown in light blue. Front, side, and top views are shown. **D)** Front view, overlay of preinitiation complex TFIIE (light coral) (PDB 7NVU) and RNA polymerase II-DSIF-ELOF1 structural model. RNA polymerase II shown as a gray surface, ELOF1 in light green, SPT5's NGN domain in dark green, template DNA in blue, non-template DNA in cyan. **E)** Top view, overlay as in D). NELF-A tentacle is colored in orange and SPT4 in light green. **F), G)** Cryo-EM density (colored surface) of template DNA (blue)-RNA (red) hybrid in the sidetracked state. Bridge helix is shown in green, metal A in magenta and nontemplate DNA in cyan. F) from DSIF-NELF-ELOF1 composite-map in the posttranslocated state and G) in the sidetracked state from DSIF-ELOF1 sidetracked map. **H)** Bacterial elemental pause sequence scaffold used for RNA extension assays[58,59]. **I)** RNA primer extension time course experiments with 10 μM CTP and GTP and in the absence or presence of indicated elongation factors. All experiments were performed at least three times, and a representative gel is shown. **J)** Intensity ratio of extended (+7) band relative to the RNA primer quantified from I).
